# Neural correlates of flexible sound perception in the auditory midbrain and thalamus

**DOI:** 10.1101/2024.04.12.589266

**Authors:** Rose Ying, Daniel J. Stolzberg, Melissa L. Caras

**Author notes:** **Correspondence:** Rose Ying, Biology-Psychology Building 4094 Campus Dr, College Park, MD 20742. **Author contributions:** M.L.C and R.Y. designed the research; R.Y. performed the research; R.Y. and D.J.S. coded and performed the analysis; R.Y. and M.L.C. wrote the paper.

## Abstract

Hearing is an active process in which listeners must detect and identify sounds, segregate and discriminate stimulus features, and extract their behavioral relevance. Adaptive changes in sound detection can emerge rapidly, during sudden shifts in acoustic or environmental context, or more slowly as a result of practice. Although we know that context- and learning-dependent changes in the spectral and temporal sensitivity of auditory cortical neurons support many aspects of flexible listening, the contribution of subcortical auditory regions to this process is less understood. Here, we recorded single- and multi-unit activity from the central nucleus of the inferior colliculus (ICC) and the ventral subdivision of the medial geniculate nucleus (MGV) of Mongolian gerbils under two different behavioral contexts: as animals performed an amplitude modulation (AM) detection task and as they were passively exposed to AM sounds. Using a signal detection framework to estimate neurometric sensitivity, we found that neural thresholds in both regions improved during task performance, and this improvement was driven by changes in firing rate rather than phase locking. We also found that ICC and MGV neurometric thresholds improved and correlated with behavioral performance as animals learn to detect small AM depths during a multi-day perceptual training paradigm. Finally, we reveal that in the MGV, but not the ICC, context-dependent enhancements in AM sensitivity grow stronger during perceptual training, mirroring prior observations in the auditory cortex. Together, our results suggest that the auditory midbrain and thalamus contribute to flexible sound processing and perception over rapid and slow timescales.

**Significance statement:** What a listener hears depends on several factors, such as whether the listener is attentive or distracted, and whether the sound is meaningful or irrelevant. Practice can also shape hearing by improving the detection of particular sound features, as occurs during language or musical learning. Understanding how changes in sound perception are implemented in the brain is important for developing strategies to optimize healthy hearing, and for treating disorders in which these processes go awry. We report that neurons in auditory midbrain and thalamus exhibit rapid shifts in sound sensitivity that depend on the sound’s behavioral relevance, and slower improvements that emerge over several days of training. Our results suggest that subcortical areas make an important contribution to flexible hearing.

## Introduction

Sound perception depends on the behavioral state and past experience of the listener. Enhanced detection or discrimination abilities can emerge rapidly due to sudden shifts in attention or expectations (Hoskin et al., 2014; Huyck & Johnsrude, 2012; Jaramillo & Zador, 2010; Kaya & Elhilali, 2017; Lawrance et al., 2014; Shinn-Cunningham, 2008; Snyder et al., 2012) and more slowly, over the course of perceptual learning (Bradlow et al., 1997; Fitzgerald & Wright, 2011; Fu & Galvin, 2007; Karawani et al., 2015; Strange & Dittmann, 1984; Wright & Sabin, 2007). In both cases, perceptual improvements are accompanied by, and often correlate with, changes in the responsiveness or tuning of auditory cortical neurons (Alain et al., 2007; Bao et al., 2004; Beitel et al., 2003; Caras & Sanes, 2017, 2019; Carcea et al., 2017; David et al., 2012; Fritz et al., 2003; Lee & Middlebrooks, 2011; Miller et al., 1972; Niwa et al., 2012; Polley et al., 2006; Recanzone et al., 1993; Rutkowski & Weinberger, 2005; van Wassenhove & Nagarajan, 2007; von Trapp et al., 2016; Whitton et al., 2014; Witte & Kipke, 2005).

Although the contribution of the auditory cortex to perceptual flexibility is well-established, anatomical and physiological findings suggest that subcortical auditory areas may also play an important role. Both the inferior colliculus (IC) and the medial geniculate nucleus (MGN) receive inputs from neuromodulatory and limbic structures that are well-positioned to mediate context-dependent and/or learning-related plasticity (Fitzpatrick et al., 1989; Klepper & Herbert, 1991; Liu et al., 2021; Marsh et al., 2002; Moore et al., 1978; Motts & Schofield, 2009, 2010; Nevue et al., 2016; Olaźbal & Moore, 1989). IC and MGN neurons also exhibit rapid changes in activity when animals perform sound-guided tasks (Franceschi & Barkat, 2021; Metzger et al., 2006; Otazu et al., 2009; Rocchi & Ramachandran, 2020; Ryan et al., 1984; Ryan & Miller, 1977, 1978; Shaheen et al., 2021; Slee & David, 2015; von Kriegstein et al., 2008) and longer-lasting changes induced by classical conditioning and associative learning (Edeline & Weinberger, 1991; Gao & Suga, 1998, 2000; Gilad et al., 2020; Lennartz & Weinberger, 1992; Taylor et al., 2021).

One limitation of the existing literature is the reliance on simple, suprathreshold sounds to assess plasticity in the IC and MGN. Few studies have used complex or near-threshold stimuli, where contextual cues or experience would provide the most benefit to the listener. Those that did use near-threshold sounds did not report neural measures of signal detectability or discriminability, instead focusing exclusively on changes in firing rate (Ryan et al., 1984; Ryan & Miller, 1977). This approach is insufficient, as indiscriminate changes in response gain have little to no impact on neurometric performance (for example, see Shaheen et al. 2021). As a result, context- and learning-dependent changes in IC and MGN sound *sensitivity* remain largely unexplored.

Here, we recorded neural activity from the central nucleus of the IC (ICC) and the ventral subdivision of the MGN (MGV) during passive sound exposure and as animals trained on an amplitude modulation (AM) detection task. Using a signal detection framework to estimate neurometric sensitivity, we found that ICC and MGV neurons exhibit better AM depth thresholds during task performance compared to passive sound exposure sessions, and that this context-dependent shift is driven by changes in firing rate rather than phase locking. We also show that neurometric thresholds in both the ICC and MGV improve as animals learn to detect smaller AM depths over several days of training, and neurometric thresholds predict behavioral performance. Finally, we reveal that in the MGV, but not the ICC, context-dependent enhancements in AM sensitivity grow stronger over the course of perceptual training. Together, these findings indicate that the IC and MGN support both context-dependent and learning-related perceptual flexibility, raising the possibility that similar changes in cortical sensitivity are inherited from the ascending auditory pathway.

## Materials and methods

### Animals

Mongolian gerbils (*Meriones unguiculatus*) were obtained from Charles River and bred in-house. Gerbils were group-housed on a 12-hour light cycle and given *ad libitum* access to chow (Purina Mills Lab Diet 5001). During behavioral training, animals were maintained on controlled water access at no less than 80% of their initial weight. To account for any effects of sex, we used approximately equal numbers of males and females. All procedures were conducted at the University of Maryland College Park and were approved by the University of Maryland Institutional Animal Care and Use Committee.

### Electrophysiology components and assembly

Chronic 64-channel electrode arrays (Buzsaki64_5x12-H64LP_30mm (N = 9) and A4x16-Poly2-5mm-20s-150-160-H64LP_30mm (N = 2), NeuroNexus) were attached to the footplate of a custom-made microdrive using superglue. A small protective wall was built around the connector using dental cement (Palacos). Prior to surgery, the assembled electrode was placed in an ultraviolet sterilization chamber (254 nm, UV Clave™) for 15 minutes.

### Surgical procedure

Electrode implantation was performed after animals had successfully completed the *associative learning* phase of training (see *Behavioral training and analysis*). One day prior to surgery, animals were given meloxicam (1.5 mg/kg, 1.5 mg/mL) either via oral suspension or subcutaneous injection as a preventative analgesic. On the day of surgery, animals were administered meloxicam and dexamethasone (0.35 mg/kg, 0.5 mg/mL) subcutaneously to prevent edema. Subjects were sedated in a small induction chamber in which isoflurane (5%) was continuously administered in O_2_ (2 L/min). After the animal was sedated, the fur on the animal’s head was shaved. The animal was then transferred to a warming pad on a stereotaxic device (Kopf) and secured in place with ear bars and a bite bar. A nose cone delivered a steady flow of isoflurane (1.5-2.5%) and oxygen at a flow rate of 2 L/min. Once a surgical plane of anesthesia was achieved (as evidenced by slow and steady respiration, and the absence of a toe-pinch response) ophthalmic ointment was applied to the eyes, and the surgical area was treated with alternating applications of betadine and alcohol. The skin on the top of the surgical area was removed with a scalpel and scissors to accommodate a headcap. The skull was cleaned and dried with H_2_O_2_ and scored with a scalpel. The implantation site was marked on the skull at the appropriate medial-lateral and anterior-posterior coordinates (Table 1). A 7 mm drill bit was used to create holes in the frontal bone and right parietal bone, and four to five bone screws (0-80 thread, 3/32”) were inserted. The skull was coated with luting cement (C&B-Metabond® Quick!) except at the marked implantation site and lambda.

**Table 1.**
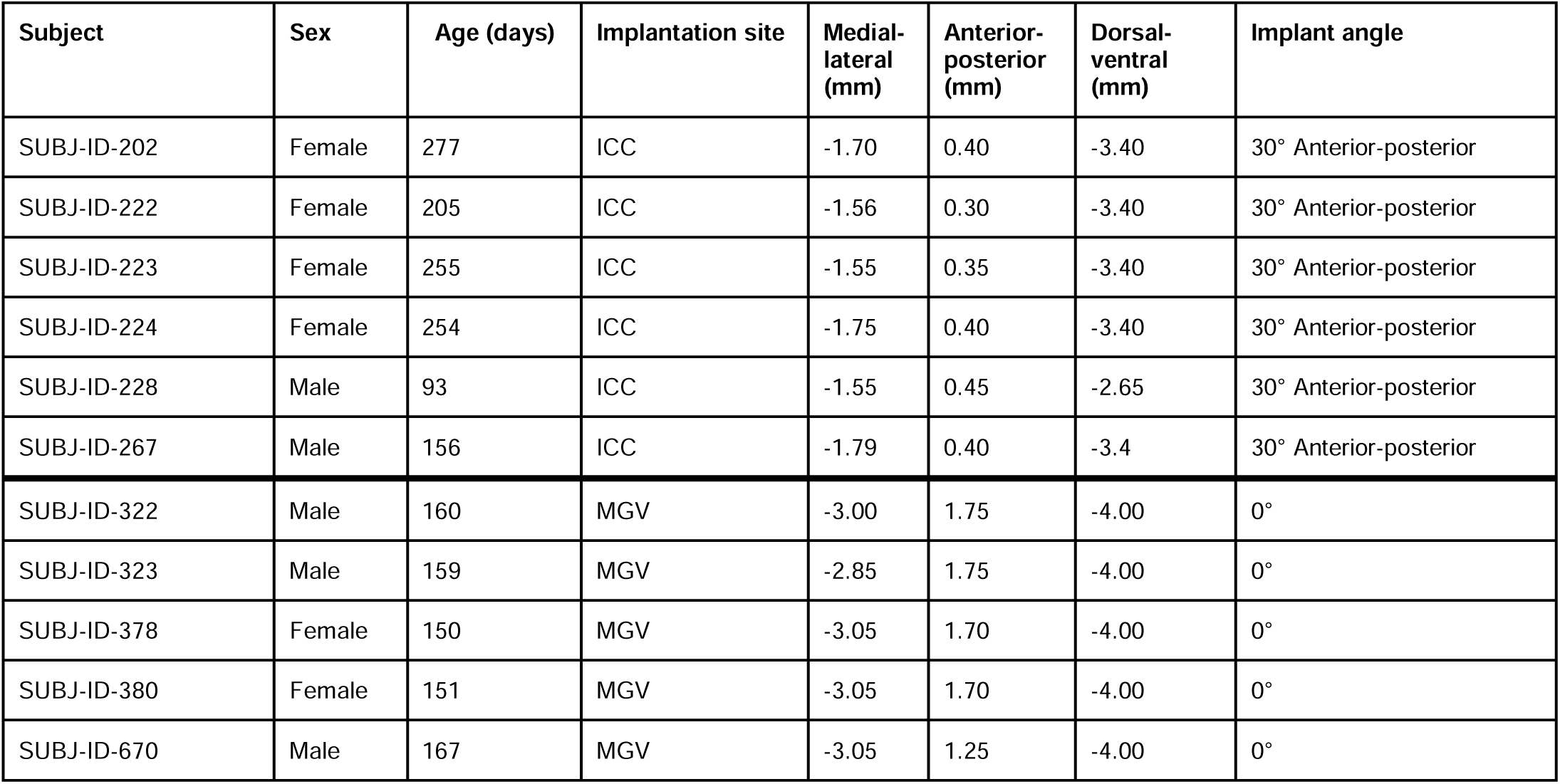
Subjects and electrode implant coordinates, relative to lambda.

A craniotomy was performed over the region of interest using a 5 mm drill bit. The assembled electrode was attached to the stereotaxic frame and fluorescent beads (0.2 µm FluoSpheres, ThermoFisher) were gently painted on the shanks using a paintbrush (Simmons et al., 2020). After lowering the electrode near the skull surface, a small burr hole was drilled away from the craniotomy site, where the ground wire was inserted and kept in place using super glue. A durotomy was performed and the electrode was lowered to the appropriate depth (Table 1). For ICC implants, the electrode was lowered at a 30-degree angle along the anterior-posterior plane. Any moveable parts of the electrode or microdrive that remained above the brain surface were coated with petroleum jelly. Dental cement was then applied in layers on top of the electrode and skull to secure the electrode. Immediately after the surgery, subjects were given a subcutaneous injection of Normosol (1-2 mL) and allowed to recover on a warming pad.

An additional dose of meloxicam and dexamethasone was administered 24 hours after surgery, and subjects were closely monitored after the surgical procedure. Animals were given a recovery period of at least one week before controlled water access began. At least one day prior to surgery and during the recovery period, subjects were treated with a prophylactic antibiotic (minocycline, 1.5 mg/kg) in their drinking water to prevent infection and reduce inflammatory processes in the brain around the electrode (Rennaker et al., 2007).

### Behavioral training and analysis

Behavioral performance on a go/no-go amplitude modulation (AM) detection task was assessed as previously described (Caras & Sanes, 2015, 2017, 2019; Macedo-Lima et al., 2023; Mowery et al., 2019; Figure 1A). Briefly, we trained animals to drink from a metal spout during continuous unmodulated broadband noise (100 Hz - 20 kHz, 45 dB SPL) and withdraw when the noise smoothly transitioned to AM (5 Hz, 1 sec duration). To encourage withdrawal during the AM stimulus, misses (drinking during AM noise) were paired with a mild shock (0.5 - 1.0 mA, 300 ms; H13-15, Colbourn).

**Figure 1.**
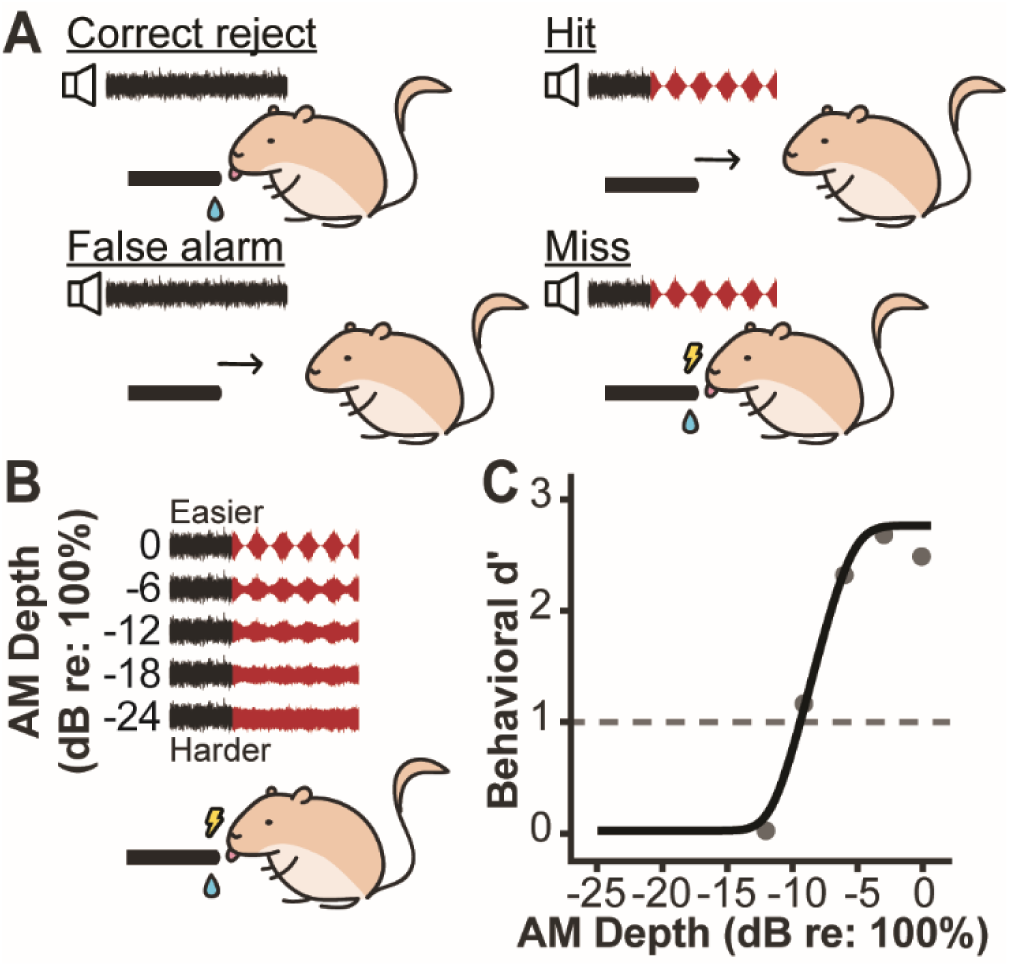
Behavioral paradigm. **A.** During *associative training*, animals learn to drink from a spout when unmodulated noise is presented, and to cease drinking when the noise transitions to AM (5 Hz, 0 dB re:100%). On unmodulated noise trials, correctly maintaining spout contact is considered a ‘correct reject’, whereas incorrectly withdrawing from the spout is considered a ‘false alarm’. On AM noise trials, correctly withdrawing from the spout is considered a ‘hit’, whereas incorrectly maintaining spout contact is considered a ‘miss’ and is punished with a mild shock. **B.** During *perceptual training*, subjects train with a wide range of AM depths. Depths gradually decrease over several days of training. Lower AM depths are harder to discriminate from unmodulated noise. **C.** Psychometric curve from a representative subject on the first day of perceptual training. Threshold is defined as the AM depth at which *d’* = 1 (dashed line).

Sounds were presented from a free-field calibrated speaker (Peerless DX25TG59-04, Tymphany) positioned 1 m above the test cage. When AM sounds were presented, the gain of the signal was adjusted to control for changes in average power across modulation depths (Viemeister, 1979; Wakefield & Viemeister, 1990). Water delivery was triggered by infrared detection of the animal at the spout and controlled by a programmable syringe pump (NE-1000, New Era). All animals were tested within a custom cage located within a Double Deluxe wooden sound booth (GretchKen) and monitored remotely via webcam. The ePsych MATLAB toolbox (Stolzberg, 2023) and custom MATLAB 2014b scripts running on a Dell PC personal computer and an RZ6 signal processor (Tucker Davis Technologies, TDT) were used to control sound output and collect behavioral data.

During the *associative training* phase, subjects learned to detect a highly salient signal (0 dB re: 100% depth AM noise). Animals first learned to drink from the spout at a 2 ml/min flow rate during 60 dB SPL unmodulated noise. Over a period of several days, the flow rate and dB SPL of the noise were lowered. Once the subject was drinking steadily at 0.4 ml/min, 0 dB re: 100% AM noise and the shock were introduced. Behavior was quantified using hits (withdrawal during AM noise) and false alarms (withdrawal during unmodulated noise) to calculate the signal detection metric *d’* (Green, 1960):

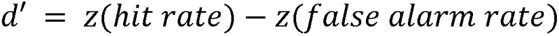

To avoid *d’* values that approach infinity, we set a floor (0.05) and a ceiling (0.95) on hit rates and false alarm rates. Once animals achieved a *d’* ≥ 2 at 45 dB SPL and 0.2 ml/min flow rate in at least one session they were surgically implanted with an electrode array. After surgery, animals were retrained on the task until they again reached a *d’* ≥ 2 at 45 dB SPL and 0.2 ml/min flow rate, after which they moved to the *perceptual training* phase.

During *perceptual training*, animals trained with a range of successively smaller AM depths for at least seven days (Figure 1B). Each day, hit rates were plotted as a function of AM depth and fit with a cumulative Gaussian using the maximum likelihood procedure of the open-source package Psignifit 4 for MATLAB (Schütt et al., 2016). In previous studies, we found that the default priors worked well for fitting similar data, so we chose to use the default priors again here (Caras & Sanes, 2015, 2017, 2019; Mowery et al., 2019). Fits were then transformed to *d’* values and used to calculate each subject’s threshold on a given day, defined as the AM depth at which *d’* = 1 (Figure 1C).

Stimuli were selected each day with the goal of bracketing the animal’s threshold, making it likely that the animal would fail to detect the smallest AM depth(s) presented. Punishing the animal by delivering a shock during these trials would likely cause the animal to either peck intermittently at the spout, resulting in an artificially high false alarm rate, or to stop approaching the spout all together. To avoid these potential confounds, the shock was turned off for the lowest two depths presented each day. A previous study tested the validity of this approach and confirmed that animals do not become conditioned to the presence or absence of the shock (Buran et al., 2014).

### Electrophysiological recording and analysis

Each day, neural recordings were made from freely moving animals during perceptual training sessions, and during periods of passive sound exposure just before (‘pre’) and just after (‘post’) the behavioral task. During passive exposure sessions, the spout was removed from the test cage, but everything else, including the sound stimuli presented and position of the recording electrodes, remained identical to the task. Animals were presented with 15 trials per AM depth during passive sessions, which closely approximated the average number of trials per depth subjects completed during the task (17.39 ± 0.3957, mean ± SEM). At the end of each training day, a subject’s electrode was advanced if there were few AM-sensitive units or if the quality of the neural signal was poor.

For some subjects (N = 5), extracellular responses were acquired with a wireless headstage and receiver (W64, Triangle Biosystems), preamplified, digitized at a 24.414 kHz sampling rate (PZ5; TDT) and fed via fiber optic link to an RZ2 BioAmp Processor (TDT) for online filtering and processing. Raw signals were streamed to an RS4 Data Streamer (TDT) for offline analysis, and data acquisition was controlled by a Dell PC running TDT’s Synapse Suite. For the remaining subjects (N = 7), responses were acquired at a 30 kHz sampling rate with a tethered RHD headstage and recording system (Intan Technologies) and a Dell PC running the Open Ephys GUI (Siegle et al., 2017).

Common average referencing and a high-pass filter (150 Hz) were applied to each channel in order to reduce noise. Spikes were extracted and sorted offline using the open-source package Kilosort2 (Pachitariu et al., 2016) (Figure 2A). The threshold for spike extraction was set to 4-10 standard deviations outside of the background noise band, and an artifact threshold was set to 50 standard deviations above the noise. Sorted spike waveforms were then clustered using principal component analysis and manually curated using phy (Rossant et al., 2023). Units were then analyzed using the Allen Brain Institute’s quality metrics (*Unit Quality Metrics*, 2023). Single units were defined as clusters with (1) a clear separation in principal component space, (2) an interspike interval (ISI) violation, which measures the rate of contaminating spikes relative to true spikes, < 0.5 (Hill et al., 2011), and (3) a fraction missing value < 0.1. In the ICC, the ISI threshold was set to 0.7 ms (Garcia-Lazaro et al., 2013; Graña et al., 2017; Yang et al., 2020). In the MGV, the ISI threshold was set to 1.5 ms. Since some MGV neurons fire in bursts, we manually examined spikes after running quality metrics for units with ISI histograms skewed towards short ISIs and a sharp decline between 3 - 10ms (Bartlett & Smith, 1999; Ramcharan et al., 2005). We classified any burst firing units which passed criteria (1) and (2) but did not pass (3) as single units, given that (3) assumes a normal amplitude histogram (Fee et al., 1996; Hill et al., 2011). Clusters that failed to meet the manual and/or quantitative criteria were considered multi-units.

**Figure 2.**
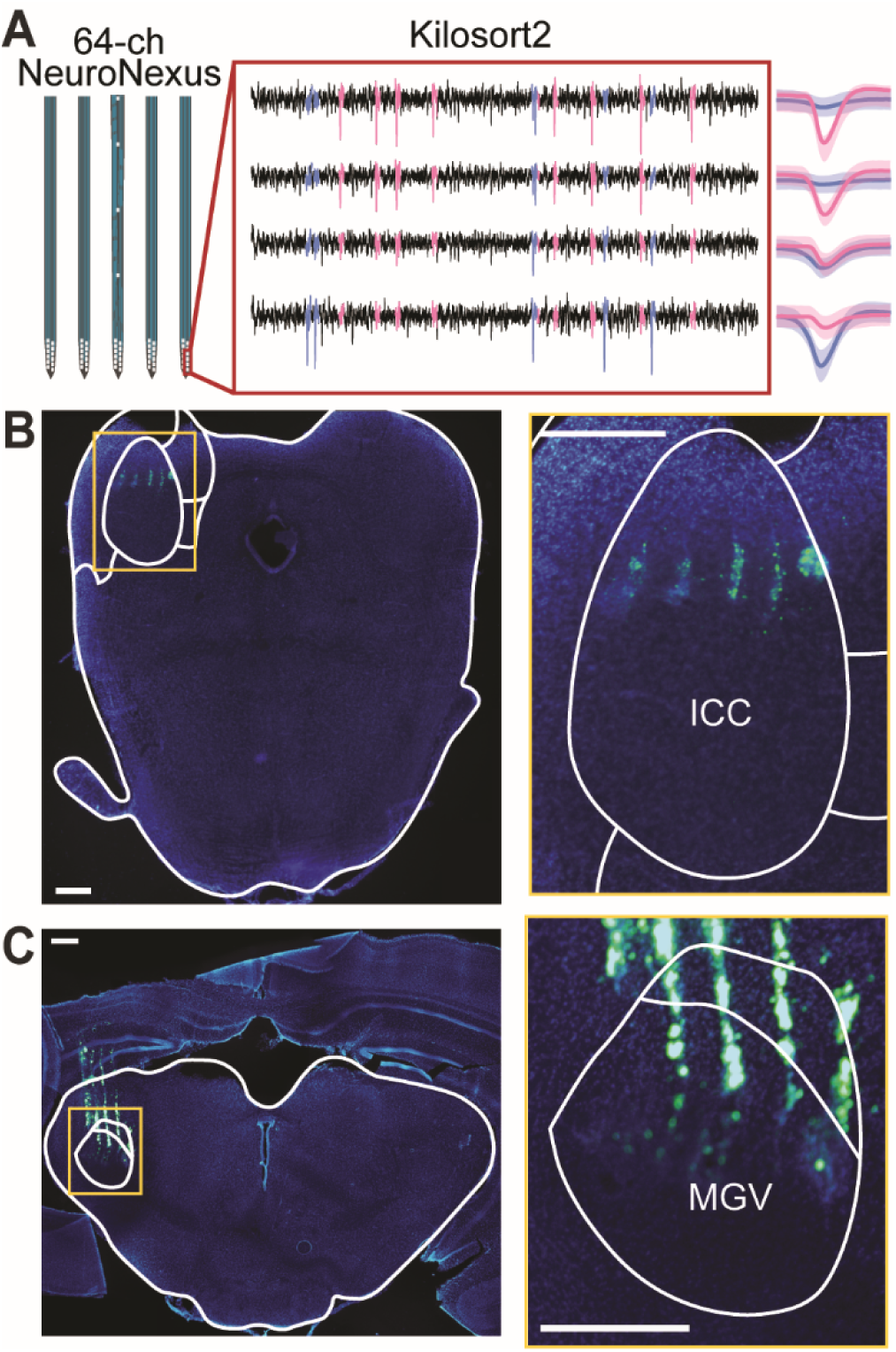
Electrophysiology. **A.** Extracellular recordings were obtained using 64-channel electrodes inserted into either ICC or MGV. Filtered voltage traces from four adjacent contact sites in the ICC are shown. Kilosort2 was used to sort spikes from individual units. **B-C.** Low and high magnification images depicting electrode tracks (green) in the ICC (**B**) and the MGV (**C**). Scale bars = 500 μm.

Each unit’s responses were quantified by using two complimentary metrics. First, each unit’s firing rate (FR; spikes/s) was calculated as the number of spikes during a one second stimulus window for both unmodulated and amplitude modulated noise. Then, each unit’s vector strength on a cycle-by-cycle basis (VScc) was calculated, as described by Yin et al., 2011. Each metric was then used to calculate a neurometric *d’* value as previously described (Caras & Sanes, 2017):

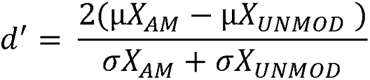

where X is the metric of interest (FR or VScc), µX_AM_ and σX_AM_ are the mean and standard deviation of X for a single modulation depth, and µX_UNMOD_ and σX_UNMOD_ are the mean and standard deviation of X during unmodulated noise. The MATLAB *nlinfit* function was used to fit neural *d’* values from a single session with a logistic function using a nonlinear least-squares regression procedure (Caras & Sanes, 2017, 2019; von Trapp et al., 2016). Neural threshold was defined as the AM depth where *d’* = 1. The validity of each fit was determined by calculating the significance of Pearson’s r correlation between predicted and actual *d’* values. Units were considered to be ‘sensitive’ to AM depth if they exhibited a significant neurometric fit (p < 0.05) and a measurable threshold (i.e., the fit crossed d’ = 1). If a unit had a significant fit, but the lowest *d’* value was above 1, threshold was defined as the smallest AM depth presented in the session. A unit was considered ‘insensitive’ to AM depth if the fit was not significant, or if the highest *d’* elicited was below 1. In many cases, units would exhibit changes in AM depth sensitivity across behavioral contexts. In these cases, unmeasurable neurometric thresholds were either assigned a value of ‘not a number’ (NaN) (Figures 4-7) or were set to 0 dB re: 100% (Figure 12) as required by the demands of the statistical analyses (see *Results* for details).

### Frequency tuning

Frequency tuning data for all units was collected on each day immediately after the post passive session. Subjects were passively exposed to a series of pure tones (500 Hz – 32 kHz in octave steps, 10-80 dB SPL in 10 dB steps, 200 msec duration, 1 sec interstimulus interval). Each frequency was played ten times in a randomized order. We then constructed frequency tuning curves by calculating the average firing rate per presented frequency. In one animal, the anatomical location of the electrode could not be conclusively determined using histology. Therefore, we used tuning curve information to determine whether the units from that animal (n = 359) displayed V-shaped narrow frequency tuning curves, characteristic of units in the MGV (Calford & Aitkin, 1983). Units that did not meet this criterion were flagged and excluded from further analysis (n = 318).

### Histology

After all training was completed, subjects were briefly placed under light isoflurane anesthesia during which electrolytic lesions were made by passing a small current (10-15 µA) through electrode channels at the tips of the shanks. Twenty-four hours later, animals were anesthetized with an intraperitoneal injection of ketamine (150 mg/kg, 25 mg/mL) and xylazine (6 mg/kg, 1 mg/mL) in saline. After the subject was determined to be insensitive to a toe pinch, the chest cavity was opened, and the subject was perfused transcardially using 1x phosphate buffered saline (PBS) followed by 4% paraformaldehyde.

### Tissue processing and microscopy

Extracted brains were post-fixed in 4% paraformaldehyde at 4°C for at least one day, then embedded in 6% agar and sliced on a vibratome (Leica VT1000 S) at 70 μm thickness. Slices were mounted on gelatin subbed slides and dried overnight before cover-slipping with ProLong Gold with DAPI. Slices were imaged on an upright fluorescent microscope (Leica DM750). To verify that recordings were made in the ICC or MGV, we confirmed that fluorescent beads, indicating electrode tracks, and/or electrolytic lesions were in our area of interest (Figure 2B-C).

### Statistical analysis

We constructed generalized linear mixed models (GLMM) in R using the glmmTMB package (Brooks et al., 2017). Normal distribution of residuals was tested after running the GLMM using DHARMa (Hartig & Lohse, 2022); if the residuals were not normal, a corrective transformation was applied to the data. An analysis of variance (ANOVA) was used to analyze the statistical significance of the GLMM. Post-hoc tests consisted of pairwise comparisons with t- and z-tests using the emmeans package (Lenth et al., 2024). All AM depth thresholds in dB were transformed back to linear percent values prior to fitting models. Results were considered statistically significant when p < 0.05. Any p values from multiple comparisons were adjusted with a Bonferroni correction.

To examine the effect of behavioral context on neurometric thresholds, firing rate ratios, and coefficients of variation, we created a model in which *Context* (i.e. pre, task, or post) was a fixed effect, and random effects included *Unit* nested within *Subject*, and *Context* nested within *Day*, using a gaussian distribution with log link. In one case, when looking at FR-based thresholds in the MGV, we unnested *Context* and *Day* to simplify the model. Due to the fact that most neurons did not exhibit a measurable FR-based threshold during pre and post sessions, FR thresholds were additionally analyzed using a binomial model with a probit link, which tested for the absence or presence of a response, with the same random effects as the gaussian model.

To determine the effect of perceptual training on behavioral thresholds, we created a model in which *Day* was a fixed effect and *Sex* nested within *Subject* was a categorical random effect, using a gaussian distribution with a log link.

To determine the effect of perceptual training on neurometric thresholds we used a model in which *Day* was a fixed effect and random effects included *Unit* nested within *Subject* and *Context* nested within *Day*, using a gaussian distribution (except for the analysis of VScc-based thresholds in the MGV, for which we used a log link).

To determine whether *Context* and *Day* had any joint effects on neurometric thresholds, we created a model in which the *Context x Day* interaction was a fixed effect and categorical random effects included *Unit* nested within *Subject* and *Session* nested within *Day*. In this analysis, any unmeasurable thresholds were replaced with a value of 0. We modeled the data using a zero-inflated beta family model, and additionally transformed linearized thresholds into a true zero-inflated distribution by using the absolute value of the linear thresholds minus 1.

To determine the correlation between behavior and neural thresholds, we used Spearman’s rank correlation coefficient.

Finally, to analyze the change in threshold from pre to task contexts we used the Rayleigh test of uniformity from the circular statistics package (Lund et al., 2023).

## Results

To characterize rapid, context-dependent shifts in sound sensitivity in the auditory midbrain and thalamus, we trained Mongolian gerbils on an aversive go/no-go amplitude modulation (AM) detection task (Caras & Sanes, 2017, 2019; Macedo-Lima et al., 2023; Mowery et al., 2019; von Trapp et al., 2016). All animals (N = 11 across all experiments) learned the task quickly, reaching our predetermined criterion level of performance (*d’* ≥ 2) within seven days (Figure 3). High levels of performance were maintained or quickly restored following surgical implantation of an electrode array into the central nucleus of the inferior colliculus (ICC, Figure 3A) or the ventral subdivision of the medial geniculate nucleus (MGV, Figure 3B). We then recorded neural responses from a total of 650 units (456 single-unit and 194 multi-units) as animals trained with a range of successively smaller AM depths over the course of several days. Each day, recordings were made while animals performed the task and during passive sound exposure sessions just before (pre) and just after (post) the task.

**Figure 3.**
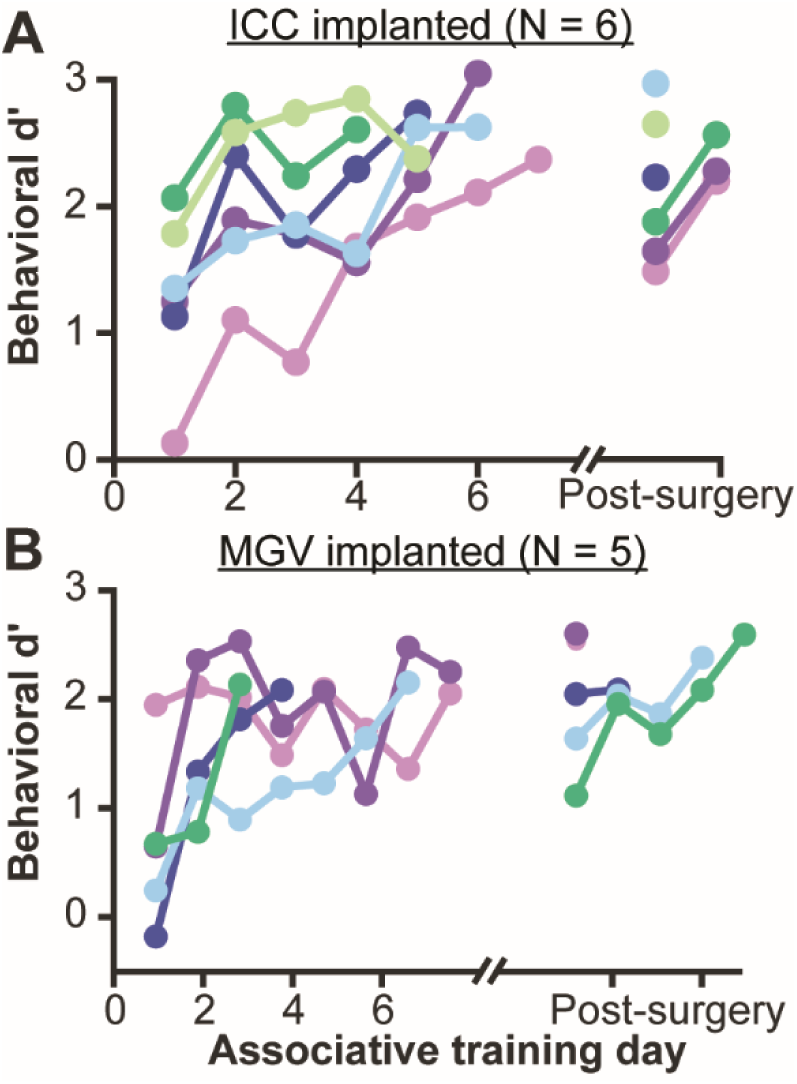
All animals successfully learned to report the presence of 0 dB re:100% depth AM noise. **A-B.** All animals reached our predetermined criterion level of performance (*d’* ≥ 2) within seven days of associative training. High performance levels were maintained or quickly restored after electrode implantation into the ICC (**A**) or MGV (**B**). Within each panel, each subject is represented by a different color.

### I Rapid shifts in spike rate, but not timing, mediate context-dependent improvements in AM sensitivity

#### I.1 Task performance improves the FR-based sensitivity of ICC and MGV neurons

Prior work has revealed that sound-evoked firing rates in both the ICC and the MGV depend on behavioral context (Franceschi & Barkat, 2021; Ryan & Miller, 1977; Saderi et al., 2021; Slee & David, 2015). While this work has yielded important insights, we do not fully understand whether or how context-dependent changes in firing rate translate into changes in the ability of subcortical neurons to detect or discriminate behaviorally relevant sounds. To answer this question, we first examined how behavioral performance on an AM detection task affects the firing rate (FR)-based AM sensitivity of ICC neurons by converting FRs into *d’* values, fitting the data, and estimating neurometric thresholds from the fits (see *Materials and Methods* for full details). For this analysis, we pooled data from units across seven days of passive sound exposure and task performance to maximize our statistical power. Units were considered ‘sensitive’ to AM depth if we could successfully estimate a neurometric threshold, and ‘insensitive’ if we could not (see *Materials and Methods* for details).

In the ICC we recorded activity from 329 total (217 single-unit, 112 multi-units). Although the majority of these units (288/329, 88%) were always insensitive to AM, a small population (41/329, 12%) was sensitive under at least one behavioral context. Data from one of these units are depicted in Figure 4A-B. During passive sound exposure sessions, AM sensitivity was extremely poor, with FR-based *d’* values remaining below 1 and resulting in unmeasurable (NaN) neurometric thresholds. In contrast, AM sensitivity was better during task performance: *d’* values increased and a neurometric threshold was measurable. This neuron was therefore considered to be ‘insensitive’ to AM during passive sound exposure, and ‘sensitive’ during task performance.

**Figure 4.**
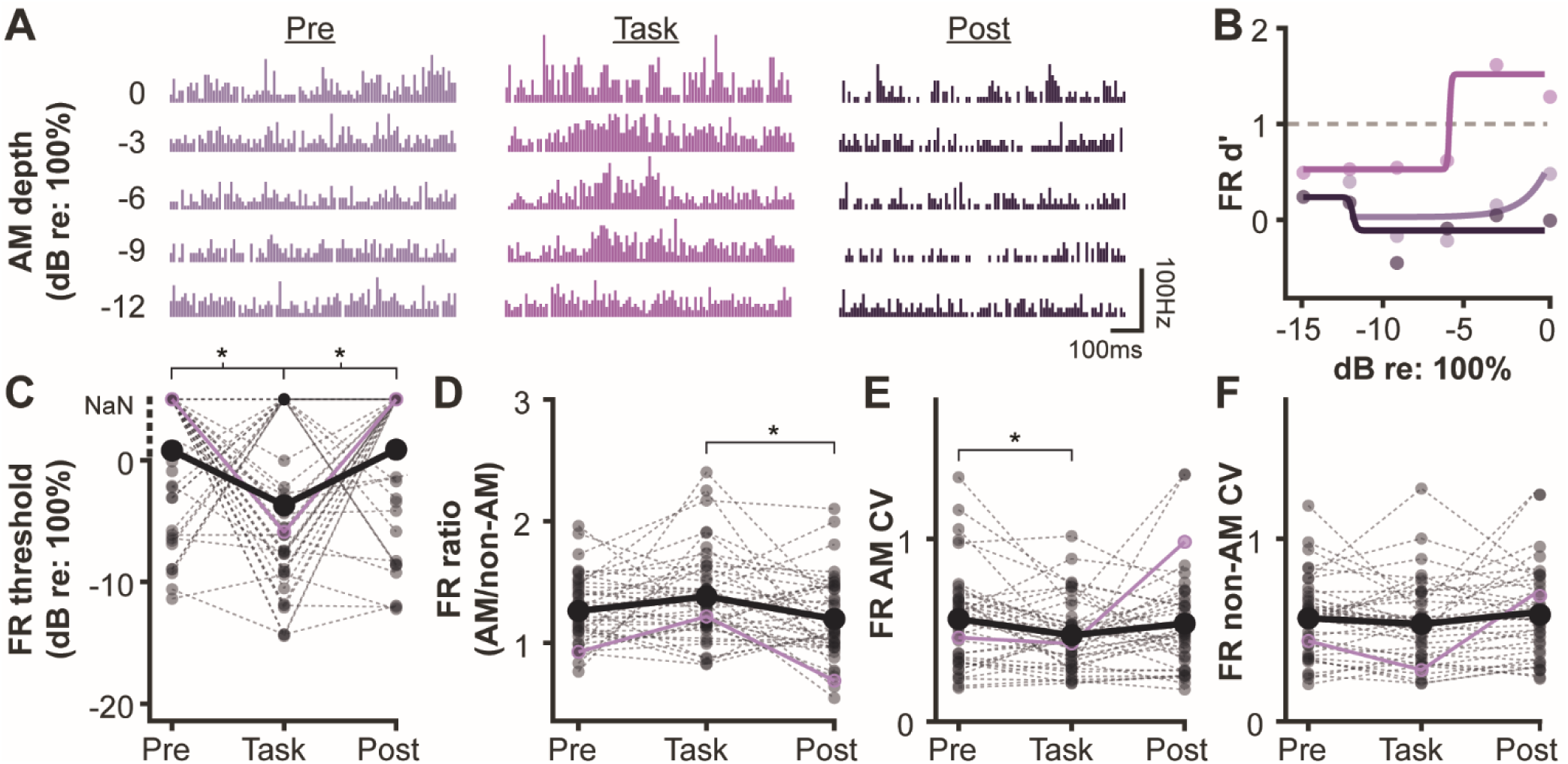
Task performance improves the FR-based sensitivity of ICC neurons. **A.** Peristimulus time histograms for one representative ICC neuron under different behavioral contexts. **B.** Neurometric functions for the same neuron depicted in A. **C.** FR-based thresholds for all ICC units that were AM sensitive in at least one behavioral context. When thresholds were unmeasurable, they were assigned a value of ‘Not a Number’ (NaN; see *Materials and Methods*). Representative neuron depicted in A and B is highlighted in purple. Dashed lines connect thresholds from the same unit. Thick black lines connect mean threshold values of all units across contexts, with each NaN replaced with a value of 5 for the calculation. **D-F.** FR ratio (**D**), coefficient of variation (CV) during AM stimulus presentation (**E**), and CV during non-AM stimulus presentation (**F**) for all ICC units that were AM sensitive in at least one behavioral context. Plot conventions are the same as panel C. * p < 0.05

To determine whether this pattern was consistent across the ICC population, we first used a binomial GLMM/ANOVA model to ask whether AM-sensitive units were more likely to exhibit a measurable threshold under a specific behavioral context. We found a significant effect of context on measurability (χ^2^(2) = 8.086, p = 0.0175, Figure 4C). AM-sensitive units were more likely to exhibit measurable FR-based thresholds during task performance than during the pre (z = 2.477, p = 0.0398) or post (z = 2.789, p = 0.0159) passive sound exposure sessions.

We then used a gaussian GLMM/ANOVA to ask whether, when measurable, threshold values differed across behavioral contexts. Again, we found a significant effect (χ^2^(2) = 25.458, p = 2.964e-06, Figure 4C). AM-sensitive ICC units exhibited lower (better) FR-based thresholds during the task (t_50_ = 4.005, p = 0.0006) and post passive session (t_50_ = 3.329, p = 0.0049) than during the pre passive exposure session. There was no significant difference between task and post sessions (t_50_ = 1.403, p = 0.5006).

What drives these context-dependent changes in FR-based sensitivity? One explanation is that task performance increases the separation of AM- and non-AM-evoked firing rate distributions, leading to higher *d’* values and lower thresholds. To explore this possibility, we calculated the ratio of average firing rates (AM/non-AM) for each ICC unit that exhibited a measurable AM threshold under at least one behavioral context. For this analysis, we focused on responses to -9 dB re: 100% depth, an AM stimulus that was presented across all days to all subjects. As shown in Figure 4D, FR ratios were modestly, but significantly affected by behavioral context (χ^2^(2) = 7.2851, p = 0.0262). FR ratios were elevated during task performance compared to the post (t_115_ = 2.601, p = 0.0315) passive sound exposure session. There was no significant difference between pre (t_115_ = 1.824, p = 0.2123) and task sessions.

Another possible driver of increased sensitivity during task performance is a reduction firing rate variability. We therefore calculated coefficient of variation (CV) values for each ICC unit that exhibited a measurable AM threshold under at least one behavioral context, again using responses to the -9 dB re: 100% depth stimulus. As shown in Figure 4E, there was a small, but significant effect of behavioral context on AM-evoked CV values (χ^2^(2) = 9.1175, p = 0.0105), such that CV values were lower during task performance than during the pre (t_115_ = -2.944, p = 0.0118) passive sound exposure session. Task CV values trended towards but were not significantly lower than the post CV values (t_115_ = -2.247, p = 0.0796). Changes in non-AM-evoked CV values were not significant across behavioral contexts (χ^2^(2) = 3.1315, p = 0.2089, Figure 4F).

We next examined how task performance affects the FR-based AM sensitivity of MGV neurons, again pooling data from across seven days of passive sound exposure and task performance. In general, we found the same pattern of results as reported for the ICC. Of 321 total recorded MGV units (239 single-unit, 82 multi-units), 117 (∼36%) exhibited a measurable FR-based threshold in at least one behavioral context. Data from a representative MGV neuron are shown in Figure 5A-B. This neuron exhibited poor AM sensitivity during passive sound exposure sessions but improved dramatically during task performance. To determine whether this pattern was consistent across the MGV population, we again used a binomial GLMM/ANOVA to ask whether AM-sensitive units were more likely to exhibit a measurable threshold under a specific behavioral context. As illustrated in Figure 5C, we found a significant effect of context (χ^2^(2) = 63.278, p = 1.817e-14), such that units were far more likely to exhibit measurable thresholds during task performance than during the pre (z = 28.954, p < 0.0001) or post (z = 32.374, p < 0.0001) passive sound exposure sessions. We then ran a gaussian GLM/ANOVA, which further revealed that when measurable, threshold values were affected by context (χ^2^(2) = 13.774, p = 0.0010), with thresholds significantly lower (better) during task performance than during the pre (t_130_ = -2.608, p = 0.0305) or post (t_130_ = -2.905, p = 0.0130) passive sound exposure sessions.

**Figure 5.**
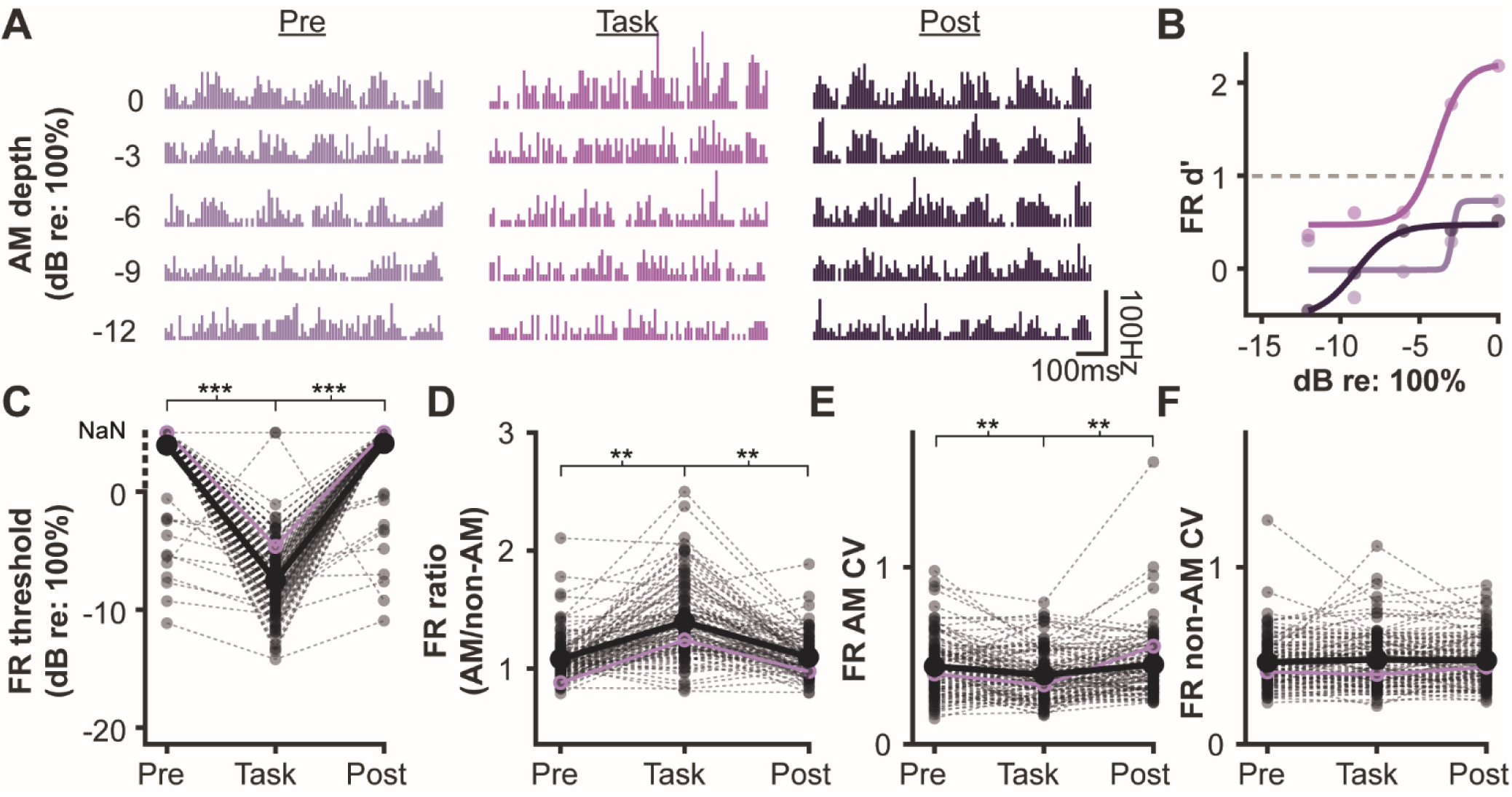
Task performance improves the FR-based sensitivity of MGV neurons. **A.** Peristimulus time histograms for one representative MGV neuron under different behavioral contexts. **B.** Neurometric functions for the same neuron depicted in A. **C.** FR-based thresholds for all MGV units that were AM sensitive in at least one behavioral context. When thresholds were unmeasurable, they were assigned a value of ‘Not a Number’ (NaN; see *Materials and Methods*). Representative neuron depicted in A and B is highlighted in purple. Dashed lines connect thresholds from the same unit. Thick black lines connect mean threshold values of all units across contexts with each NaN replaced with a value of 5 for the calculation. **D-F.** FR ratio (**D**), coefficient of variation (CV) during AM stimulus presentation (**E**), and CV during non-AM stimulus presentation (**F**) for all MGV units that were AM sensitive in at least one behavioral context. ** p < 0.01, *** p < 0.0001.

To examine if the enhanced AM sensitivity during task performance was driven by an increase in the separation between AM- and non-AM firing rate distributions, we calculated the ratio of average firing rates (AM/non-AM) for each individual MGV neuron that exhibited a measurable AM threshold under at least one behavioral context. As above, we restricted this analysis to -9 dB re: 100% depth. MGV FR ratios were significantly affected by behavioral context (χ^2^(2) = 79.505, p = 2.2e-16, Figure 5D). FR ratios during task performance were significantly higher than those during the pre (t_343_ = 8.027, p < 0.0001) and post (t_343_ = 7.425, p < 0.0001) passive sound exposure sessions.

We then analyzed CV values for each MGV unit that exhibited a measurable AM threshold under at least one behavioral context to determine whether changes in firing rate variability also contribute to context-dependent shifts in MGV sensitivity. During presentation of the -9 dB re:100% AM stimulus, there was a significant effect of context on CV values (χ^2^(2) = 17.52, p = 0.0002; Figure 5E). Firing variability was significantly lower during task performance than during both pre (t_343_ = -3.057, p = 0.0072) and post (t_343_ = -3.753, p = 0.0006) passive sessions. In contrast, context did not affect firing variability during non-AM noise (χ^2^(2) = 0.54, p = 0.7627; Figure 5F).

For all of the above analyses, similar findings were observed if we restricted our analysis to just single units (see Table 2 for statistics; data not shown). Collectively, these results suggest that in ICC and MGV neurons, task performance drives changes in both the magnitude and variability of spiking, leading to enhanced detectability of the target AM sound.

**Table 2.** Effect of context on FR-based metrics for single units.

| Region | Test | Result | p-value |
| --- | --- | --- | --- |
| ICC | Threshold (sensitive/not sensitive) | $\chi^2(2) = 14.225$ | 0.0008 |
| | Task-Pre | $z = 3.289$ | 0.0030 |
| | Task-Post | $z = 4.037$ | 0.0002 |
| | Measurable thresholds | $\chi^2(2) = 5.9024$ | 0.0529 |
| | Task-Pre | $t_{28} = -2.043$ | 0.1517 |
| | Task-Post | $t_{28} = -0.185$ | 1.0000 |
| | AM/non-AM ratio | $\chi^2(2) = 11.367$ | 0.0034 |
| | Task-Pre | $t_{73} = 2.557$ | 0.0379 |
| | Task-Post | $t_{73} = 3.100$ | 0.0083 |
| | AM CV | $\chi^2(2) = 8.229$ | 0.0163 |
| | Task-Pre | $t_{73} = -1.986$ | 0.1525 |
| | Task-Post | $t_{73} = -2.808$ | 0.0192 |
| | Non-AM CV | $\chi^2(2) = 2.0602$ | 0.3570 |
| MGV | Threshold (sensitive/not sensitive) | $\chi^2(2) = 51.816$ | 5.6e-12 |
| | Task-Pre | $z = 22.829$ | < 0.0001 |
| | Task-Post | $z = 20.675$ | < 0.0001 |
| | Measurable thresholds | $\chi^2(2) = 13.529$ | 0.0012 |
| | Task-Pre | $t_{93} = -2.594$ | 0.0331 |
| | Task-Post | $t_{93} = -2.750$ | 0.0215 |
| | AM/non-AM ratio | $\chi^2(2) = 68.503$ | 1.333e-15 |
| | Task-Pre | $t_{250} = 7.410$ | < 0.0001 |
| | Task-Post | $t_{250} = 6.916$ | < 0.0001 |
| | AM CV | $\chi^2(2) = 16.499$ | 0.0003 |
| | Task-Pre | $t_{250} = -3.317$ | 0.0031 |
| | Task-Post | $t_{250} = -3.692$ | 0.0008 |
| | Non-AM CV | $\chi^2(2) = 0.3855$ | 0.8247 |

#### I.2 Behavioral context has a heterogeneous effect on the VScc-based sensitivity of ICC and MGV neurons

In addition to firing rate, context-dependent changes in spike timing can also contribute to enhanced sound sensitivity (Niwa et al., 2012; Sameiro-Barbosa & Geiser, 2016). We therefore asked whether task performance affects the AM sensitivity of ICC or MGV neurons when estimated using a measure of neural phase-locking by calculating the cycle-by-cycle vector strength (VScc) for each stimulus and each unit. We then converted VScc values into *d’* values, fit the data, and estimated neurometric thresholds from the fits (see *Materials and Methods* for full details). As above, we pooled data from across days to maximize our statistical power. Units were considered ‘sensitive’ to AM depth if we could successfully estimate a neurometric threshold, and ‘insensitive’ if we could not (see *Materials and Methods* for details). For these analyses, we only included single units.

When we examined VScc-based sensitivity in the ICC, over half of the single-units (120/217, 55%) were always insensitive to AM depth. The remainder (n = 97/217; 45%) were sensitive during at least one behavioral context. In general, these neurons followed one of three distinct response patterns. A substantial portion (46/97, ∼47%) were sensitive during the task and during at least one passive sound exposure session. Data from one of these neurons are depicted in Figure 6A-B. This neuron was not only always sensitive to AM, but its sensitivity remained fairly constant across all behavioral contexts. This observation was generally true for the other neurons that fell into this subcategory; there was no significant effect of behavioral context on their VScc-based thresholds (χ^2^(2) = 3.2299, p = 0.1989, Figure 6C). The remaining two groups of neurons were affected by behavioral context, but in opposing manners. One group of cells (39/97, ∼40%) were AM-sensitive during passive sound exposure sessions but insensitive during the task (Figure 6D-E). A final group of neurons (12/97, ∼12%) were AM insensitive during passive sound exposure sessions but AM sensitive during the task (Figure 6F-G). As a result of this response heterogeneity, average VScc-based thresholds across the ICC population did not significantly differ across the pre, task, or post conditions (χ^2^(2) = 1.3429, p = 0.511, Figure 6H).

**Figure 6.**
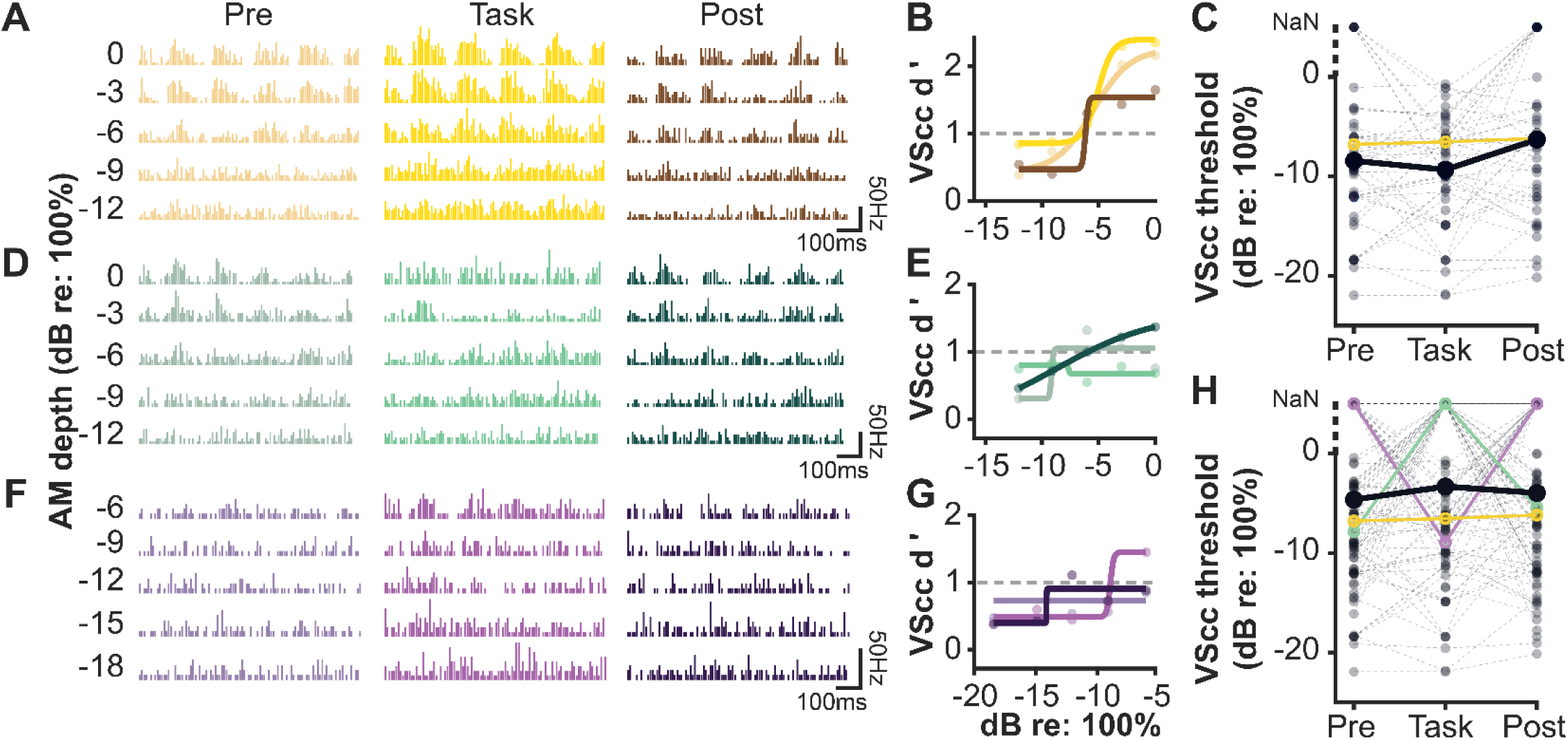
Behavioral context has a heterogenous effect on the VScc-based sensitivity of ICC neurons. **A.** PSTH for one representative ICC neuron sensitive to AM depth under all behavioral contexts **B.** Neurometric functions for the same neuron as depicted in A. **C.** VScc-based thresholds for all ICC neurons sensitive to AM depth during task performance and at least one passive session. When thresholds were unmeasurable, they were assigned a value of ‘Not a Number’ (NaN; see *Materials and Methods*). Dashed lines connect thresholds from the same unit. Thick black lines connect population means for each context with each NaN replaced with a value of 5 for the calculation. Representative neuron depicted in A and B is highlighted in yellow. **D.** PSTH for one representative ICC neuron sensitive to AM depth only during passive sound exposure sessions. **E.** Neural *d’* values and fits for the same neuron as depicted in D. **F.** PSTH for one representative ICC neuron sensitive to AM depth only during the task. **G.** Neural *d’* values and fits for the same neuron as depicted in F. **H.** VScc-based thresholds for all ICC neurons that were AM sensitive in at least one behavioral context. Dashed lines connect thresholds from the same unit. Thick black lines connect population means for each context with each NaN replaced with a value of 5 for the calculation. Representative neurons are highlighted in the same colors as in previous panels.

We next looked at how task performance affects the VScc-based AM sensitivity of MGV neurons, again pooling data as in our previous analysis. A minority of sampled MGV neurons (n = 91/606, 15%) were AM sensitive under at least one behavioral context, and as in the ICC, three distinct response patterns were observed. Of the AM-sensitive MGV neurons, 37/91 (40%) were sensitive during at least one passive session and during task performance. Data from one of these neurons are depicted in Figure 7A-B. On average, the sensitivity of these neurons did not significantly change across contexts (χ^2^(2) = 2.9737, p = 0.2261; Figure 7C). A second group of neurons (36/91, 40%) were AM sensitive during at least one passive sound exposure session but insensitive during the task (Figure 7D-E). The final group (18/91, 20%) were insensitive to AM during passive sessions but sensitive during the task (Figure 7F-G). Across the entire population, average VScc-based thresholds did not significantly differ across the pre, task, or post conditions (χ^2^(2) = 4.3168, p = 0.1155, Figure 7H). Together, these findings suggest that in both ICC and MGV neurons, task performance does not drive a systematic shift in spike-timing-based AM detection, and instead generates heterogenous changes across the neuronal population.

**Figure 7.**
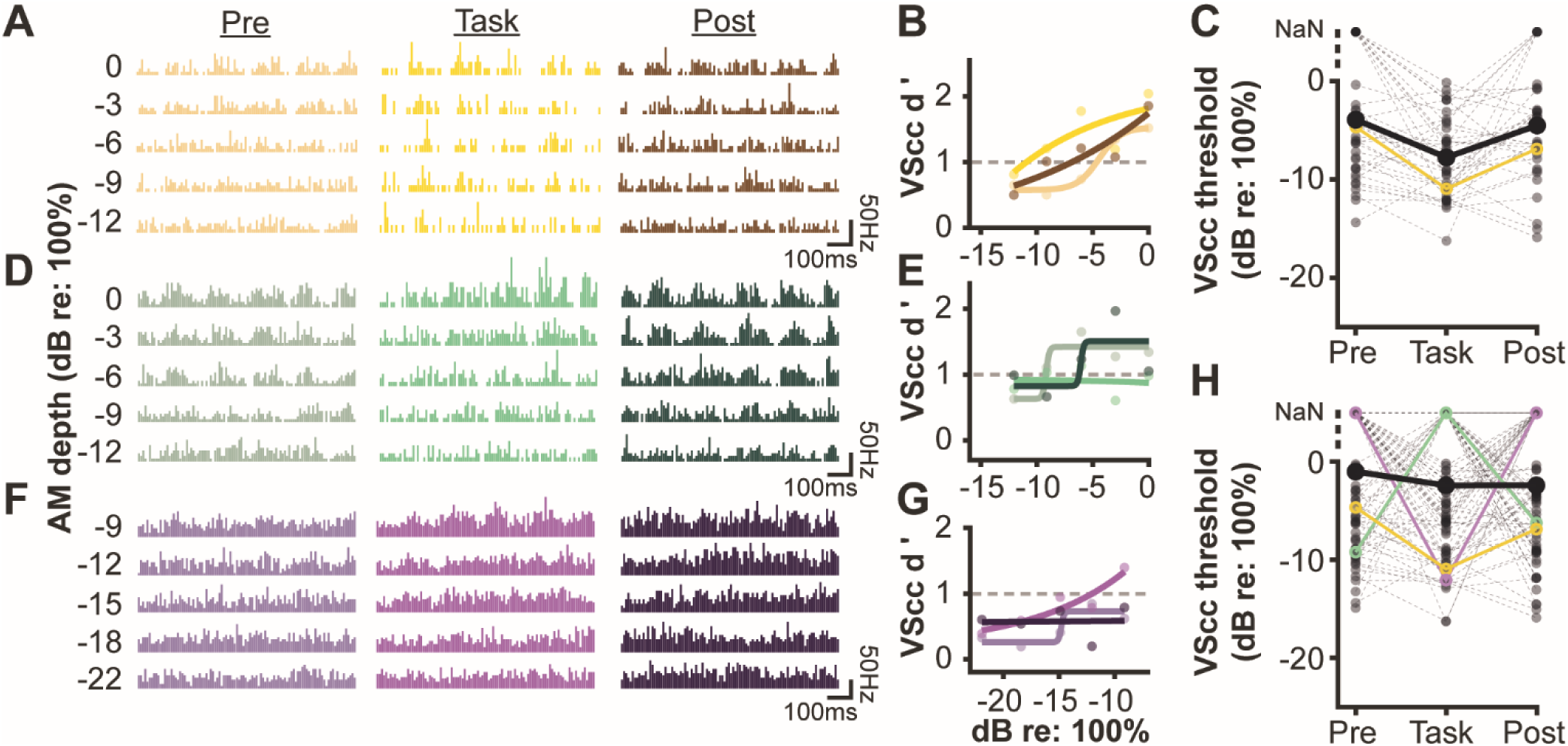
Behavioral context has a heterogeneous effect on the VScc-based sensitivity of MGV neurons. **A.** PSTH for one representative MGV neuron sensitive to AM depth under all behavioral contexts. **B.** Neurometric functions for the same neuron as depicted in A. **C.** VScc-based thresholds for all MGV neurons sensitive to AM depth during task performance and at least one passive session. When thresholds were unmeasurable, they were assigned a value of ‘Not a Number’ (NaN; see *Materials and Methods*). Dashed lines connect thresholds from the same unit. Thick black lines connect population means for each context with each NaN replaced with a value of 5 for the calculation. Representative neuron depicted in A and B is highlighted in yellow. **D.** PSTH for one representative MGV neuron sensitive to AM depth only during passive sound exposure sessions. **E.** Neural *d’* values and fits for the same neuron as depicted in D**. F.** PSTH for one representative MGV neuron sensitive to AM depth only during the task. **G.** Neural *d’* values and fits for the same neuron as depicted in F. **H.** VScc-based thresholds for all MGV neurons that were AM sensitive in at least one behavioral context. Dashed lines connect thresholds from the same unit. Thick black lines connect population means for each context with each NaN replaced with a value of 5 for the calculation. Representative neurons are highlighted in the same colors as in previous panels.

#### I.3 Context-dependent shifts in FR-based and VScc-based thresholds largely occur independently

To gain a better understanding of the relationship between VScc-based and FR-based shifts in sensitivity within individual neurons, we created single-unit population vector plots for each brain region (Figure 8A-B). In each plot, context-dependent shifts were calculated as the difference between thresholds obtained during the task and thresholds obtained during the pre passive sound exposure session. For this analysis, all single units that exhibited a measurable FR-based or VScc-based AM threshold during at least one of these sessions were included, and unmeasurable thresholds (NaNs) were set to a value of 0 dB re: 100%. Each vector (grey arrows) represents the direction and magnitude of the FR-based and VScc-based shifts for an individual neuron.

**Figure 8.**
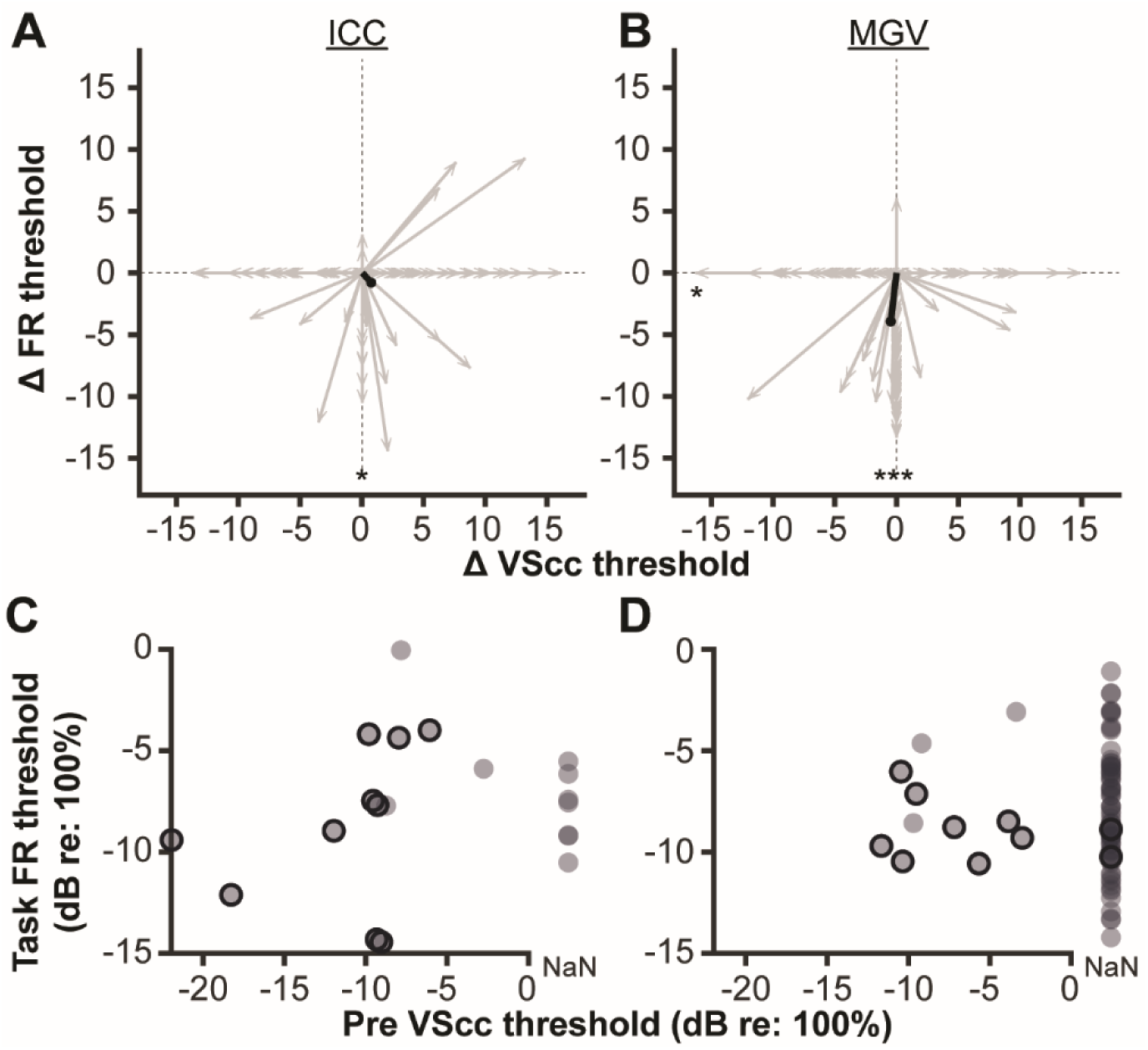
Context-dependent shifts in FR-based and VScc-based thresholds are largely independent. **A.** Each vector (grey arrow) represents the direction and magnitude of the VScc-based and FR-based thresholds shifts for an individual ICC neuron. Shifts were calculated as the difference between thresholds obtained during the task and thresholds obtained during the pre passive sound exposure session. Negative shifts reflect instances when the threshold was lower (better) during task performance. Black line represents vector mean. **B.** Same as panel A, but for MGV neurons. **C.** FR-based thresholds from ICC neurons obtained during task performance are plotted against VScc-based thresholds obtained during the pre passive sound exposure session. Each circle represents data from one single-unit. When VScc-based pre passive thresholds were unmeasurable, they were assigned a value of ‘Not a Number’ and plotted separately to the right of the y axis. Neurons with a measurable VScc task threshold are outlined in black. **D.** Same as panel C, but for MGV neurons. *p < 0.05, ***p < 0.0001

In the ICC, 76/92 (∼83%) neurons shifted their AM sensitivity along a single dimension, exhibiting a context-dependent change in their FR-based or VScc-based threshold, but not both. These neurons fall along the x and y axes in Figure 8A. A smaller subset, 16/92 (∼17%), changed along both dimensions. Overall, there was no systematic relationship between the direction of the VScc-based and FR-based shifts; while most of these neurons improved their FR-based threshold from pre to task sessions (Rayleigh test statistic: 0.1311, p = 0.0376), they were equally likely to improve or worsen along the VScc-based dimension (Rayleigh test statistic: -0.0977, p = 0.9047).

Similar results were observed in the MGV. Of 144 neurons, 128 (91%) shifted their AM sensitivity along a single dimension, and the remainder (13/144, 9%) shifted along both (Figure 8B). Nearly all neurons that shifted their FR-based sensitivity improved during the task (Rayleigh test statistic: 0.5604, p = 1.860e-21). VScc-based AM sensitivity was also slightly more likely to improve (Rayleigh test statistic: 0.1320, p = 0.0125). Together, these data suggest that the mechanisms that drive context-dependent shifts in subcortical spike rate and spike timing operate largely independently.

#### I.4 In a subset of neurons, AM detection is supported by changes in both firing rate and phase locking during task performance

Many ICC and MGV neurons exhibited a measurable FR-based threshold exclusively during task performance (Figure 4C and 5C). This observation raised two possibilities. First, these neurons may be largely insensitive to AM depth during passive exposure, and only detect AM sounds when they gain behavioral relevance. Alternatively, these neurons may in fact detect AM stimuli during passive sound exposure, but do so by phase locking, rather than increasing their firing rate. To distinguish these possibilities, we compared FR-based thresholds obtained during task performance with VScc-based thresholds obtained during the pre passive sound exposure period. For this analysis, we only included single units that exhibited a measurable FR-based threshold during the task, and as above, we pooled data across days to increase our statistical power.

As illustrated in Figure 8C, roughly half of the ICC neurons that exhibited a measurable FR-based threshold during the task (13/20) exhibited a measurable VScc-based threshold during the pre passive sound exposure session. Of these neurons, the majority (10/13, ∼79%) also had measurable VScc-based thresholds during task performance (indicated by the outlined circles in Figure 8C). In the MGV, a much smaller proportion of neurons that exhibited a measurable FR-based threshold during the task exhibited a measurable VScc-based threshold during passive sound exposure (11/84, ∼13%, Figure 8D). Of those that did, we found that a majority (8/11, ∼73%) also exhibited measurable VScc-based thresholds during task performance. Together, these findings suggest that while some neurons, particularly those in the MGV, are only sensitive to AM depth when AM sounds are behaviorally relevant, others are always sensitive, but switch from relying solely on phase-locking during passive sound exposure to using a combination of phase-locking and overall spike rate when performing a task. Moreover, these data indicate that the reliance on a rate-based AM detection strategy increases as information moves up the ascending auditory pathway, consistent with previous literature (Bartlett & Wang, 2007; Cariani, 1999; Joris et al., 2004; Joris & Yin, 1992; Krishna & Semple, 2000; Wang, 2007; Wang et al., 2008).

### II Subcortical sensitivity to sound gradually improves across perceptual learning

During perceptual training, subjects improve their ability to detect or discriminate increasingly difficult stimulus cues over several days or weeks of practice. These perceptual enhancements are accompanied by training-induced changes in stimulus representations within the auditory cortex (Alain et al., 2007; Caras & Sanes, 2017, 2019; Polley et al., 2006; Recanzone et al., 1993; van Wassenhove & Nagarajan, 2007; Witte & Kipke, 2005). Whether neurons in subcortical auditory structures undergo similar changes remains uncertain. To address this issue, we estimated the AM sensitivity of ICC and MGV neurons as animals learned to detect smaller and smaller AM depths.

#### II.1 Perceptual training improves behavioral AM depth sensitivity

We first assessed behavioral AM detection thresholds across seven days of perceptual training with a range of AM depths. Figure 9A-B depicts data from a representative animal, illustrating improved psychometric performance across training. As expected from prior work (Caras & Sanes, 2015, 2017, 2019; Mowery et al., 2019), behavioral thresholds significantly decreased across training days in animals used for ICC (χ^2^(6) = 45.834, p = 3.195e-08, Figure 9C) and MGV (χ^2^(6) = 21.570, p = 0.0014, Figure 9D) recordings. These data confirm that animals underwent perceptual learning on our AM depth detection task.

**Figure 9.**
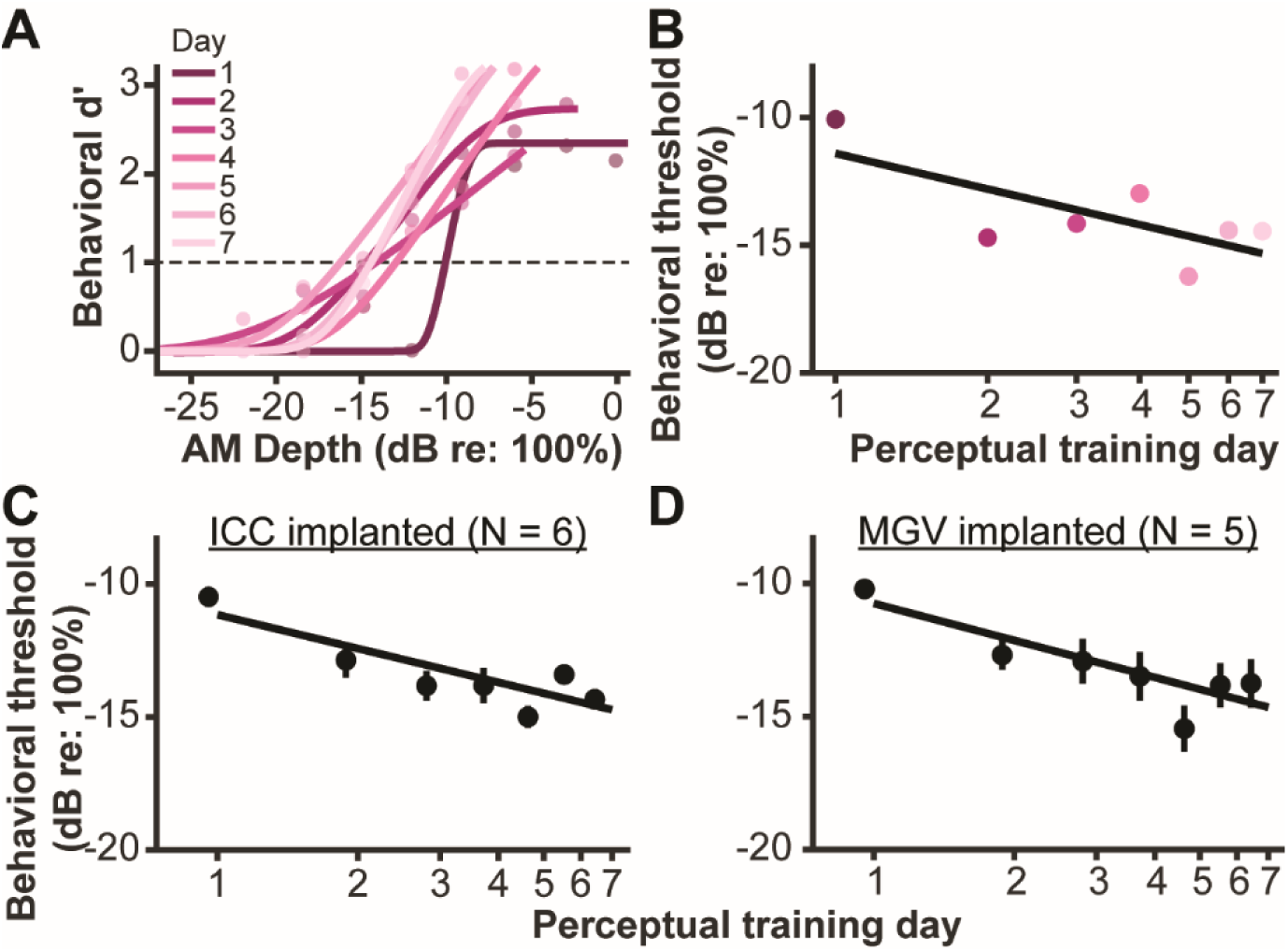
Perceptual training improves behavioral AM depth thresholds. **A.** Psychometric fits from one representative animal across seven days of perceptual training. **B.** Behavioral thresholds from the same animal depicted in A. **C-D.** Mean +/- SEM behavioral thresholds for all animals implanted with electrodes in the ICC (N = 6, **C**) or MGV (N = 5, **D**).

#### II.2 FR-based thresholds of ICC and MGV neurons improve during perceptual training and predict perceptual performance

We asked whether FR-based thresholds of ICC neurons change during perceptual training. For this analysis, we only included units that were deemed ‘sensitive’ to AM noise (i.e. exhibited a measurable FR-based threshold) during the task. Because relatively few ICC single-units met this criterion (20 neurons across all animals and days), we pooled data from single and multi-units to increase our statistical power. Figure 10A reveals that when averaged across all units and animals, FR-based thresholds significantly decrease (improve) over the course of perceptual training (χ^2^(5) = 13.025, p = 0.0232). Neural and behavioral improvement occurred in tandem, with the average sensitivity of ICC neurons closely tracking average perceptual sensitivity across training days. Therefore, we asked whether average neural thresholds predicted behavioral thresholds in individual animals. Data from one representative subject is presented in Figure 10B. We did not observe a significant correlation between FR-based ICC thresholds and behavioral thresholds within this subject (r = 0.7629, p = 0.2371), likely because the data were underpowered. However, when we pooled data across all subjects, a significant correlation was observed (r = 0.6451 p = 0.0029, Figure 10C).

**Figure 10.**
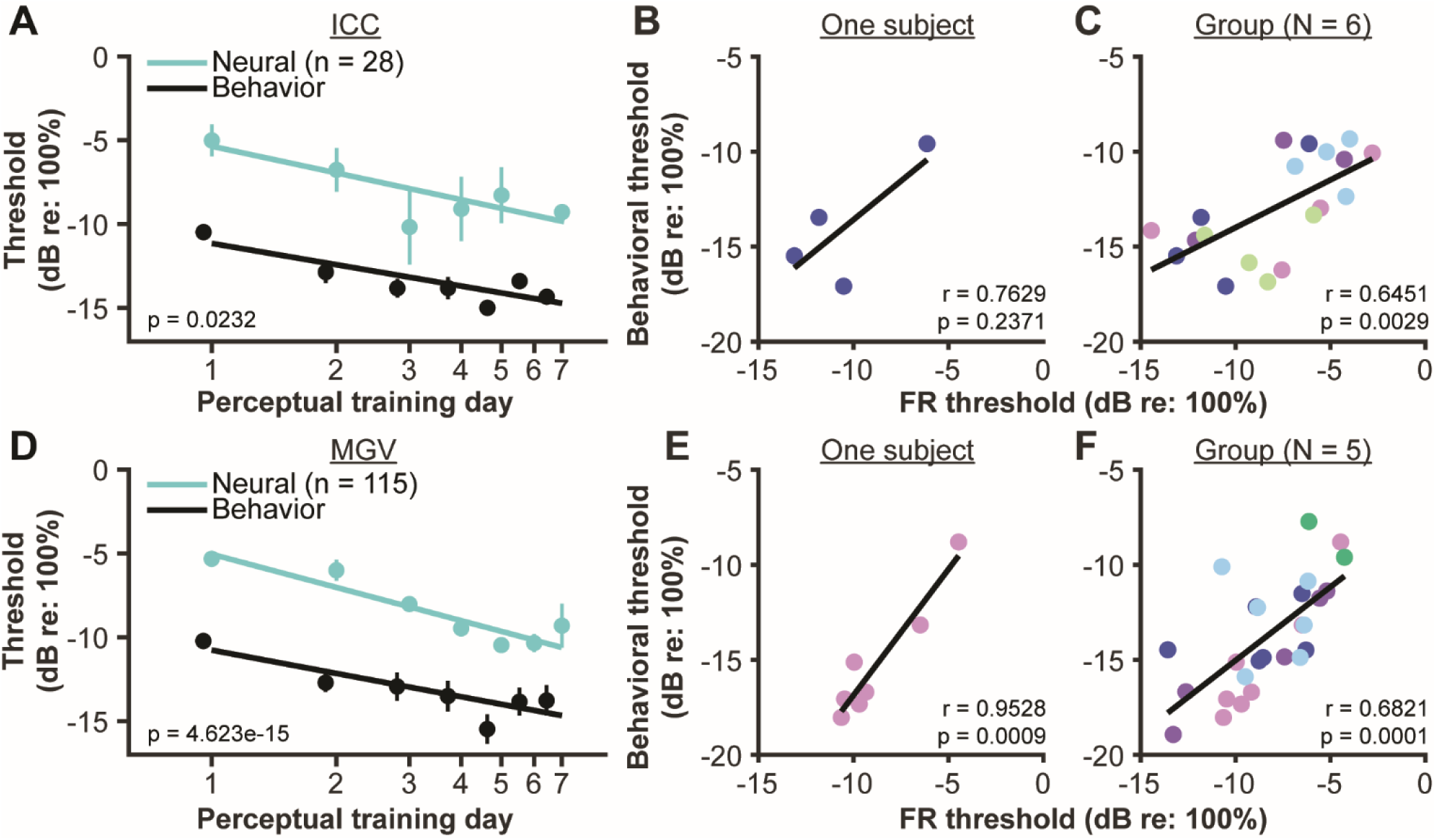
FR-based thresholds of ICC and MGV neurons improve during perceptual training and predict perceptual performance. **A.** Mean +/- SEM FR-based ICC thresholds decrease across perceptual training. Thresholds were estimated from AM-sensitive units during task performance (n = 28 units; 2 - 7/day). Behavioral thresholds from Figure 9 are re-plotted here for comparison with neural data. **B.** Representative data from one subject. Each data point represents the average FR-based ICC threshold obtained during task performance for a given training day, plotted against the animal’s behavioral threshold for the same day. **C.** FR-based neural thresholds in the ICC significantly correlate with behavioral thresholds. Each color represents data from an individual animal. **D.** Mean +/- SEM FR-based MGV thresholds decrease across perceptual training. Thresholds were estimated from AM-sensitive units during task performance (n = 115 units; 2 – 31/day). Behavioral thresholds from Figure 9 are re-plotted here for comparison with neural data. **E.** Representative data from one subject. Data are as described in panel B. **F.** FR-based neural thresholds in the MGV significantly correlate with behavioral thresholds across animals. Each color represents an individual animal.

Next, we examined whether perceptual training affects the FR-based AM sensitivity of MGV neurons, again pooling data from single- and multi-units that were deemed AM-sensitive during the task. In general, we found the same pattern of results as reported for the ICC, where FR-based thresholds of MGV neurons improved over the course of training (GLMM/ANOVA, χ^2^(6) = 74.608, p = 4.623e-15; Figure 10D). The rate of improvement (-6.57 dB/log(day)) was similar to that observed in the ICC (-5.29 dB/log(day)). We then again asked whether average neural thresholds predicted behavioral thresholds in individual animals. Correlations were significant in 2/5 animals. Data from a subject exhibiting a significant correlation is depicted in Figure 10E (r = 0.9528, p = 0.0009). Neural and behavioral thresholds were also significantly correlated when we pooled data across all subjects (Figure 10F; r = 0.6821, p = 0.0001). Collectively, these results indicate that rate-based representations of AM stimuli in the ICC and MGV significantly improve and correlate with behavioral thresholds during perceptual training.

#### II.3 VScc-based thresholds of ICC and MGV neurons improve during perceptual training and predict perceptual performance

Improvements in neural phase locking to auditory stimuli may also underlie longer-term learning (Bao et al., 2004; Batterink, 2020), particularly in subcortical regions, where temporal representations of sound stimuli are more prevalent (Bartlett & Wang, 2007; Joris et al., 2004; Krishna & Semple, 2000; Wang et al., 2008). To determine whether this is the case during perceptual learning, we explored whether VScc-based thresholds improved across days of perceptual training. We restricted this analysis to single units that were AM-sensitive during the task.

In the ICC, average VScc-based thresholds appeared to improve with training (-3.48 dB/log(day)), but the effect did not reach statistical significance (χ^2^(6) = 11.665, p = 0.0699, Figure 11A). VScc-based thresholds only correlated with behavioral thresholds in 1 of 5 subjects. In the other 4 subjects, there was no correlation, as illustrated for one representative animal in Figure 11B (r = 0.2597, p = 0.6731). At the group level, there was a moderate correlation between neural and behavioral thresholds (r = 0.4361, p = 0.0203; Figure 11C).

**Figure 11.**
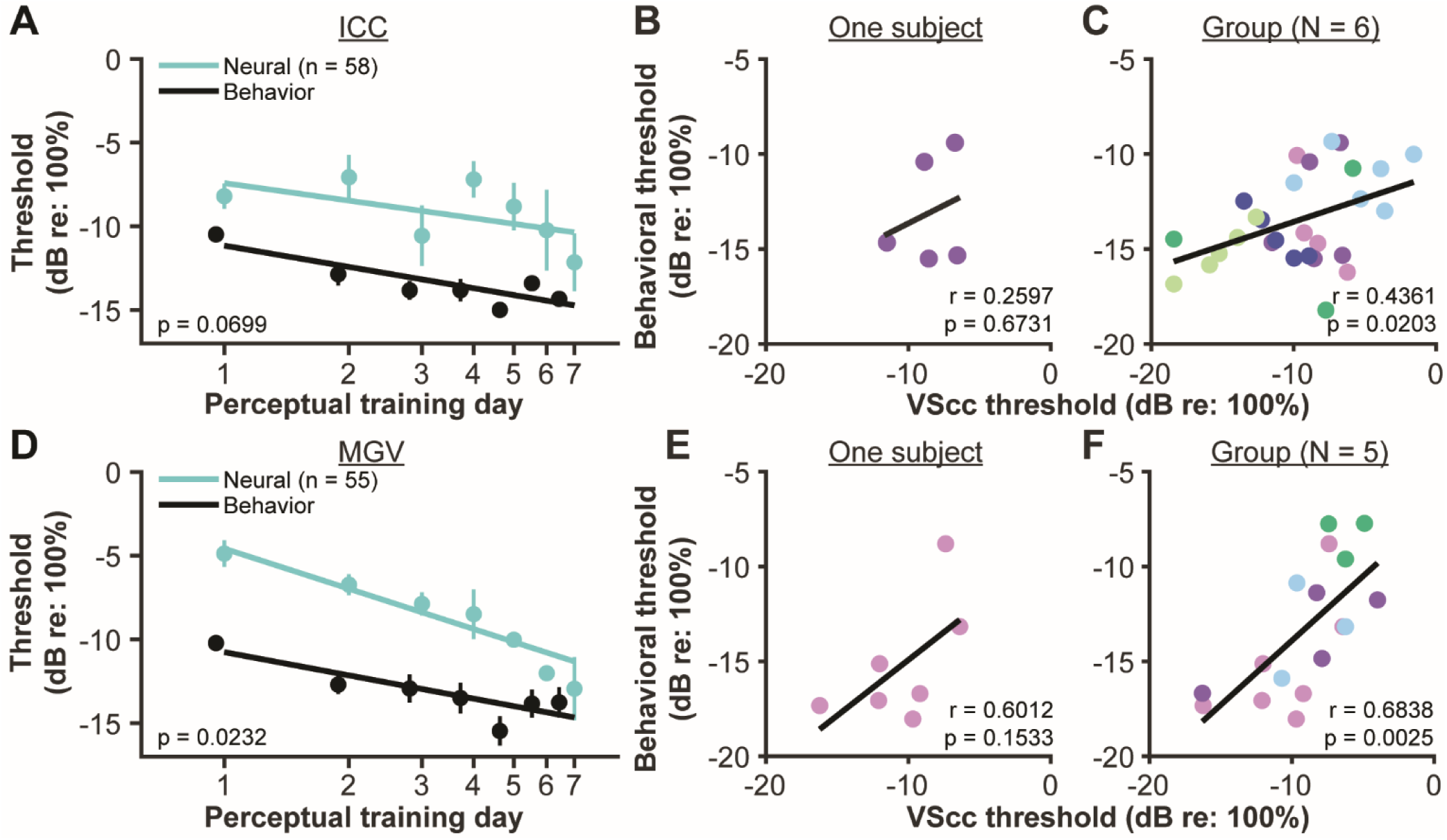
VScc-based thresholds of ICC and MGV neurons improve during perceptual training and predict perceptual performance. **A.** Mean +/- SEM VScc-based ICC thresholds decrease across perceptual training. Thresholds were estimated from AM-sensitive units during task performance (n = 58 units; 3 - 14/day). Behavioral thresholds from Figure 9 are re-plotted here for comparison with neural data. **B.** Representative data from one subject. Each data point represents the average VScc-based ICC threshold obtained during task performance for a given training day, plotted against the animal’s behavioral threshold for the same day. **C.** VScc-based neural thresholds in the ICC significantly correlate with behavioral thresholds across animals. Each color represents an individual animal. **D.** Mean +/- SEM VScc-based MGV thresholds decrease across perceptual training. Thresholds were estimated from AM-sensitive units during task performance (n = 55 units; 1 - 19/day). Behavioral thresholds from Figure 9 are re-plotted here for comparison with neural data. **E.** Representative data from one subject. Data are as described in panel B. **F.** VScc-based neural thresholds in the MGV significantly correlate with behavioral thresholds across animals. Each color represents an individual animal.

We then looked at whether training had any effect on VScc-based thresholds in the MGV. In contrast to the ICC, VScc-based thresholds significantly improved across training (GLMM/ANOVA, χ^2^(6) = 26.049, p = 0.0002; Figure 11D). However, VScc-based MGV thresholds did not correlate with behavioral thresholds among any of the individual subjects, as illustrated for a representative animal in Figure 11E (r = 0.6012, p = 0.1533). Thresholds were significantly correlated at the group level, however (r = 0.6838, p = 0.0028; Figure 11F).

These data indicate that phase-locked representations of AM stimuli improve and correlate with behavioral thresholds during perceptual learning but do so more inconsistently or weakly than rate-based representations, and thus offer less explanatory power.

### III Context-dependent improvements in neural sensitivity increase during perceptual training

#### III.1 The magnitude of context-dependent effects on FR-based thresholds increases during perceptual training in the MGV, but not the ICC

Auditory cortical neurons are more sensitive to AM stimuli when subjects perform an AM detection task than during passive sound exposure, and the magnitude of this enhancement increases over the course of perceptual training (Caras & Sanes, 2017). We found that ICC and MGV units exhibit context-dependent enhancements in AM sensitivity that mirror those observed in the auditory cortex (Figures 4-5). Whether the magnitude of these context-dependent enhancements similarly increase over the course of perceptual training remains unknown.

To address this issue, we first asked whether there was a significant interaction between behavioral context and perceptual training day on FR-based thresholds in the ICC. For this analysis, we pooled data from single- and multi-units, and only included units that had at least one measurable AM threshold. Of these units, 33/41 (∼80%) had unmeasurable (‘NaN’) thresholds during the pre and/or post passive session. Excluding these units from our analysis would therefore cause us to grossly overestimate the average AM sensitivity during passive sound exposure, and therefore *underestimate* the effect of behavioral context. We therefore set these NaN values to a value of 0, the highest possible AM depth. We dropped days 4 and 6 from the analysis because all units on those days did not exhibit measurable thresholds in at least one of the three sessions leading to rank deficiency. As illustrated in Figure 12A, and as expected from the analysis presented in Figure 4C, FR-based thresholds in the ICC were strongly modulated by behavioral context (χ^2^(2) = 12.555, p = 0.0019). This effect remained relatively constant across time, however, such that there was no significant interaction between context and perceptual training day (χ^2^(8) =10.289, p = 0.2453).

**Figure 12.**
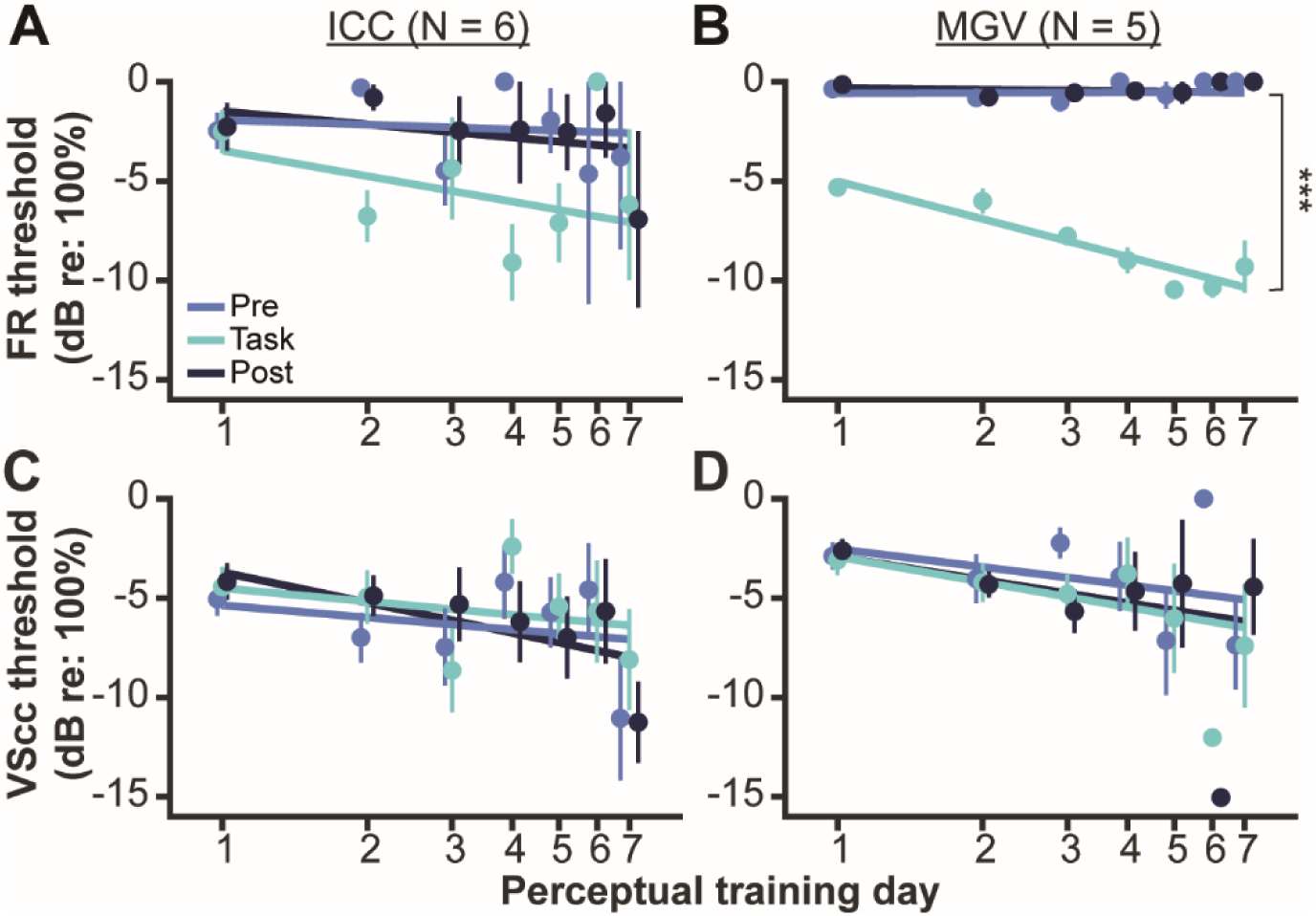
Context-dependent improvements in FR-based sensitivity increase with training in the MGV. **A.** FR-based ICC thresholds are lower (better) during task performance than passive sound exposure, but the magnitude of this context-dependent enhancement does not change across perceptual training. Data are from 41 units; 2 - 10/day. **B.** FR-based MGV thresholds are lower (better) during task performance than passive sound exposure, and the magnitude of this context-dependent enhancement increases across perceptual training. Data are from 117 units; 2 - 32/day. **C.** VScc-based ICC thresholds decrease across perceptual training but exhibit no effect of behavioral context, and no interaction between context and day. Data are from 97 units; 9 - 24/day. **D.** VScc-based MGV thresholds decrease across perceptual training but exhibit no effect of behavioral context, and no interaction between context and day. Data are from 91 units; 1 – 30/day. All data are depicted as means +/- SEMs. Thresholds were estimated from units that had at least one measurable AM threshold. NaN thresholds were replaced with a value of 0 for statistical analysis (see *Materials and Methods*). *** p < 0.0001.

We then examined FR-based thresholds in the MGV. As above, we pooled data from single- and multi-units, only included units that had at least one measurable AM threshold (n = 117) and set NaN values to 0 dB re: 100% depth to account for units with NaN thresholds in the pre and/or post session (110/117, 94%). We dropped days 4, 6, and 7 from the analysis due to rank deficiency, as described above. We again found a strong effect of behavioral context (χ^2^(2) = 46.108, p = 9.723e-11). However, as shown in Figure 12B, this effect grew stronger over time, resulting in a significant interaction between behavioral context and perceptual training day (χ^2^(6) = 23.474, p = 0.0007). These data are reminiscent of our previous observations in the auditory cortex (Caras & Sanes, 2017).

#### III.2 VScc-based ICC and MGV thresholds improve across training days regardless of context

We found that behavioral context exerts a heterogeneous influence on VScc-based thresholds in the ICC and MGV (Figures 6-7), such that there is no significant effect at the population level. This observation does not preclude the possibility, however, that the degree of heterogeneity might decrease over the course training, allowing a significant effect of context to emerge. To explore this possibility, we asked whether there was a significant interaction between behavioral context and perceptual training day on VScc-based thresholds in the ICC. We included single-units displaying a measurable VScc-based threshold in at least one of the three sessions (n = 97). Many of these neurons exhibited NaN thresholds in at least one session (95/97, 98%). Therefore, we again replaced any NaN thresholds with a value of 0. As shown in Figure 12C, we found a significant effect of day on VScc-based thresholds (χ^2^(6) = 25.944, p = 0.0002), but no effect of context (χ^2^(2) = 2.8683, p = 0.2383) and no interaction between day and context (χ^2^(12) = 7.690, p = 0.8089).

We next examined VScc-based thresholds in the MGV. A total of 91 single units had measurable VScc-based thresholds in at least one session. As in the IC, the majority had a NaN threshold in at least one of the pre, task, and post sessions (87/91, ∼96%); these unmeasurable thresholds were replaced with a value of 0. We also dropped day 6 from the model, as only one unit exhibited VScc-based sensitivity that day in the task and post sessions. The results are shown in Figure 12D and are similar to those obtained from the ICC. While VScc-based MGV thresholds significantly improved across perceptual training days (χ^2^(5) = 55.8238, p = 8.835e-11), neither the effect of context (χ^2^(2) = 2.5063, p = 0.2856) nor the interaction between day and context (χ^2^(10) = 5.3729, p = 0.8649) were significant.

Collectively, these observations suggest that the magnitude of context-dependent effects on FR-based sensitivity increases during perceptual training and arises at the level of the MGV.

## Discussion

Adaptive changes in the spectral and temporal sensitivity of auditory cortical neurons underlie many aspects of flexible listening, but the contribution of subcortical auditory stations to this process is less understood. Here we found that in the ICC and MGV, neuronal sensitivity to AM, a temporal cue that supports speech perception, is better when subjects perform an AM detection task compared to when subjects are passively exposed to AM sounds. This context-dependent effect, which is largely mediated by changes in spike rate rather than stimulus phase-locking, grows stronger over the course of perceptual training, particularly in the MGN, mirroring previous reports in the auditory cortex. These findings raise the possibility that aspects of flexible sound processing observed in the auditory cortex may actually be inherited from subcortical auditory regions.

### Effects of behavioral context on subcortical sound processing

Prior studies examining the effect of behavioral context on subcortical sound processing have yielded conflicting results. While some studies found that behavioral relevance affects IC and MGN sound-evoked responses (Franceschi & Barkat, 2021; Ryan et al., 1984; Ryan & Miller, 1977; Shaheen et al., 2021), others reported the opposite (Otazu et al., 2009; Rocchi & Ramachandran, 2020; Slee & David, 2015; Williamson et al., 2015). This confusion likely stems from several factors. First, the majority of prior studies did not specify which IC and MGN subdivisions were sampled (Franceschi & Barkat, 2021; McGinley et al., 2015; Metzger et al., 2006; Otazu et al., 2009; Ryan et al., 1984; Saderi et al., 2021). Second, the existing literature has focused almost exclusively on context-dependent changes in firing rate without considering changes in phase locking, which predominates in lower subcortical stations, particularly the IC (Joris et al., 2004). Finally, few studies have examined the effect of behavioral context on IC or MGN responses to near-threshold stimuli, and those that did stopped short of quantifying signal detectability or discriminability (Ryan et al., 1984; Ryan & Miller, 1977). Here, we targeted first-order midbrain and thalamic subdivisions and used a signal detection framework to explore how behavioral context impacts firing rate- and vector strength-based sensitivity to a stimulus cue with real-world relevance: AM depth. Our results demonstrate an unambiguous effect of context: the ICC and MGV exhibit rapid improvements in firing rate-based thresholds during task performance compared to periods of passive sound exposure. Additional experiments are needed to determine whether these results generalize to other stimuli or to reward-based tasks (David et al., 2012).

### Temporal dynamics of task-dependent changes in sound processing

Two previous studies reported that in a subset of ICC neurons, task-dependent changes in activity persist for several minutes after task performance ends (Saderi et al., 2021; Slee & David, 2015). Here, we found no evidence of such persistence in the ICC or MGV; task-dependent improvements in FR-based sensitivity generally reverted back to baseline levels immediately after task performance. This apparent discrepancy may be explained by differences in species (ferrets vs. gerbils), task structure (appetitive vs. aversive), target stimuli (pure tones vs. AM noise), and measured outcome variable (spike rate vs. neurometric threshold).

Regardless, previous work using the same species, paradigm, and stimuli as this study found that in the auditory cortex, task-dependent improvements in AM sensitivity *do* persist after the task is over (Caras & Sanes, 2017). This observation, together with our findings, suggest that separate mechanisms with distinct temporal dynamics may operate simultaneously to shape context-dependent plasticity in auditory subcortical and cortical regions. Moreover, the fact that auditory cortical persistence is often incomplete, with neural sensitivity reverting slightly, but not completely, back to pre-task levels during the post-passive period raises the possibility that context-dependent shifts in the auditory cortex may reflect a combination of rapid transient mechanisms operating subcortically, and local mechanisms operating within the auditory cortex that decay with a longer time constant.

### Models for perceptual learning

Several models have been put forth to explain perceptual learning (for review, see Dosher & Lu, 2017; Watanabe & Sasaki, 2015). One long standing model is *reverse hierarchy theory*, which suggests that plasticity first occurs within higher-order sensory brain regions, and changes in lower levels of the sensory pathway only emerge later in training (Ahissar & Hochstein, 2004). Despite this model’s prominence, it remains difficult to determine how well it describes the existing data, particularly in the auditory system. Part of this difficulty is due to the fact that relatively few studies have examined the contribution of subcortical regions to perceptual learning. Of the studies that have, all used tools that lack temporal or spatial precision, and most have restricted their analyses to time points before and after training (Carcagno & Plack, 2011; Lau et al., 2017; MacLean et al., 2024; Reetzke et al., 2018). This approach obscures the temporal relationship between training-induced changes in neural function and perception, and is problematic given that the neuroplasticity that accompanies learning is often transient, renormalizing once behavioral performance stabilizes (Floyer-Lea et al., 2006; Frank et al., 2018; Muellbacher et al., 2001; Sampaio-Baptista et al., 2015; Sarro et al., 2015; Yotsumoto et al., 2008).

We recorded from the ICC and MGV while subjects actively trained on a multi-day perceptual learning task. Three notable findings emerged. First, neuronal sensitivity to the target AM sound improved over the course of training in both regions. Second, the rate of improvement was relatively similar in the ICC and MGV and largely comparable to the rate of improvement previously observed in the auditory cortex (Caras & Sanes, 2017). Finally, neural and behavioral sensitivity improved in tandem, emerging over similar time courses and to similar degrees. Taken together, our results do not support the reverse hierarchy theory.

Another key finding from our experiments was an interaction between context-dependent and learning-dependent plasticity, such that task performance has a greater effect on rate-based AM sensitivity at the end of training than at the beginning. This effect was weak in the ICC, possibly due to a lack of statistical power, but was robust in the MGV. Similar observations in the auditory cortex previously led us to propose that non-sensory processes drive context-dependent changes in the auditory cortex, and training facilitates perceptual learning by increasing the strength of these modulations (Caras & Sanes, 2017). Our current data suggest that our observations in the auditory cortex may partially result from modulatory processes acting on subcortical regions in the ascending auditory pathway.

### Neural mechanisms driving subcortical plasticity

While we demonstrate that ICC and MGV neurons exhibit context-dependent and learning-related plasticity, the neural mechanisms underlying these changes remain uncertain. Context-dependent changes in activity have been noted as early as the auditory nerve (Delano et al., 2007; Gehmacher et al., 2022) and cochlear nuclei (Oatman, 1971; Ryan et al., 1984), and population-level activity from cochlear nucleus neurons can predict behavioral detection thresholds (Mackey et al., 2023). Plasticity in these early regions is likely mediated by descending inputs from the superior olivary complex; indeed, medial olivocochlear bundle activity increases during perceptual training on a phoneme-in-noise discrimination task (Boer & Thornton, 2008). Additional experiments in animals with silenced olivocochlear efferents will be needed to determine the extent to which our results reflect inherited changes from the auditory periphery or early brainstem nuclei.

Our data show that a behavioral shift from passive sound exposure to task engagement is accompanied by a neural shift to improved rate-based sensitivity within individual ICC and MGV neurons. While this effect is relatively small in the ICC, it increases in both significance and magnitude in the MGV, suggesting a hierarchical emergence of context-dependent plasticity which parallels the emergence of the rate code along the ascending auditory pathway. The mechanism driving the temporal-to-rate transformation is currently unknown, although recent work implicates GluN2C/D containing NMDA receptors in the IC (Drotos et al., 2023).

Other possible drivers of subcortical plasticity could involve inputs from dopaminergic, cholinergic, noradrenergic, and/or serotonergic sources (Chen et al., 2018; Fitzpatrick et al., 1989; Klepper & Herbert, 1991; Moore et al., 1978; Motts & Schofield, 2009, 2010; Nevue et al., 2016). In the ICC, for example, serotonin levels fluctuate with context (Hurley & Hall, 2011), and could mediate short-term synaptic plasticity and changes in gain (Bohorquez & Hurley, 2009; Miko & Sanes, 2009). Thus, serotonergic inputs may contribute to the rapid rate-based ICC plasticity reported here.

A final, though not mutually exclusive, possibility is that ICC and MGV plasticity is mediated by descending inputs from the auditory cortex. Layer 5 corticofugal neurons project to higher-order IC and MGN subdivisions (Asilador & Llano, 2021; Bajo & Moore, 2005; Williamson & Polley, 2019) and transmit sensory and non-sensory signals to downstream targets (Ford et al., 2024; Oberle et al., 2022). Ablation or inactivation of these neurons has a stronger impact on learning when stimuli are difficult to detect or discriminate, as in our perceptual training paradigm (Bajo et al., 2010; Ford et al., 2024; Krall et al., 2024). Coupling ICC and MGV recordings with L5 corticofugal loss-of-function manipulations will be necessary to establish whether this descending pathway makes a causal contribution to perceptual learning and its associated subcortical plasticity.

## Acknowledgements

This work was supported by National Institute of Health Grant T32DC00046 and F31DC021355 to R.Y. Purchase of the Zeiss LSM 980 Airyscan 2 was supported by Award Number 1S10OD025223-01A1 from the National Institute of Health. We thank Dr. Matheus Macedo-Lima (University of Maryland) for help with statistics and all members of the M.L.C. laboratory for their constructive criticism and support. The authors declare no competing financial interests.

